# Roles for Mitochondrial Complex I subunits in regulating synaptic transmission and growth

**DOI:** 10.1101/2021.12.30.474605

**Authors:** Bhagaban Mallik, C. Andrew Frank

## Abstract

To identify conserved components of synapse function that are also associated with human diseases, we conducted a genetic screen. We used the *Drosophila melanogaster* neuromuscular junction (NMJ) as a model. We employed RNA interference (RNAi) on selected targets and assayed synapse function and plasticity by electrophysiology. We focused our screen on genetic factors known to be conserved from human neurological or muscle functions (300 *Drosophila* lines screened). From our screen, knockdown of a Mitochondrial Complex I (MCI) subunit gene (*ND-20L*) lowered levels of NMJ neurotransmission. Due to the severity of the phenotype, we studied MCI function further. Knockdown of core MCI subunits concurrently in neurons and muscle led to impaired neurotransmission. We localized this neurotransmission function to the muscle. Pharmacology targeting MCI phenocopied the impaired neurotransmission phenotype. Finally, MCI subunit knockdowns or pharmacological inhibition led to profound cytological defects, including reduced NMJ growth and altered NMJ morphology. Mitochondria are essential for cellular bioenergetics and produce ATP through oxidative phosphorylation. Five multi-protein complexes achieve this task, and MCI is the largest. Impaired Mitochondrial Complex I subunits in humans are associated with disorders such as Parkinson’s disease, Leigh syndrome, and cardiomyopathy. Together, our data present an analysis of Complex I in the context of synapse function and plasticity. We speculate that in the context of human MCI dysfunction, similar neuronal and synaptic defects could contribute to pathogenesis.

## 1. Introduction

Mitochondrial Complex I (MCI) is a multimeric enzyme with a molecular mass of about 1MDa (Hirst, 2013). It modulates the transfer of electrons from NADH to ubiquinone, facilitating ATP synthesis (Galkin et al., 2006; Galkin et al., 1999; Wikstrom, 1984). In humans, dysfunction of MCI activity can contribute to forms of neurodegeneration, and this is thought to be due to accumulation of excess reactive oxygen species (ROS) (Reviewed by (Breuer et al., 2013)). But on neuronal and synaptic levels, how this manifests in disease is unclear.

Structurally, Mitochondrial Complex I comprises 44 subunits in mammals; 42 of those are present in *Drosophila melanogaster* MCI, which is the focus of the present study (Garcia et al., 2017; Guerrero-Castillo et al., 2017). The 42 subunits of *Drosophila* MCI are composed of 14 conserved subunits forming the catalytic core. The remaining 28 are termed accessory subunits (Garcia *et al*., 2017) (Fig. 1A-B). Even though accessory subunits are not directly involved in catalysis, prior genetic and biochemical studies of MCI indicate that disruption of accessory subunits can produce high levels of reactive oxygen species (ROS), resulting in impaired MCI assembly and stability *in vivo* (Berger et al., 2008; Formosa et al., 2020; Guerrero-Castillo *et al*., 2017; Stroud et al., 2016).

**Figure 1:**
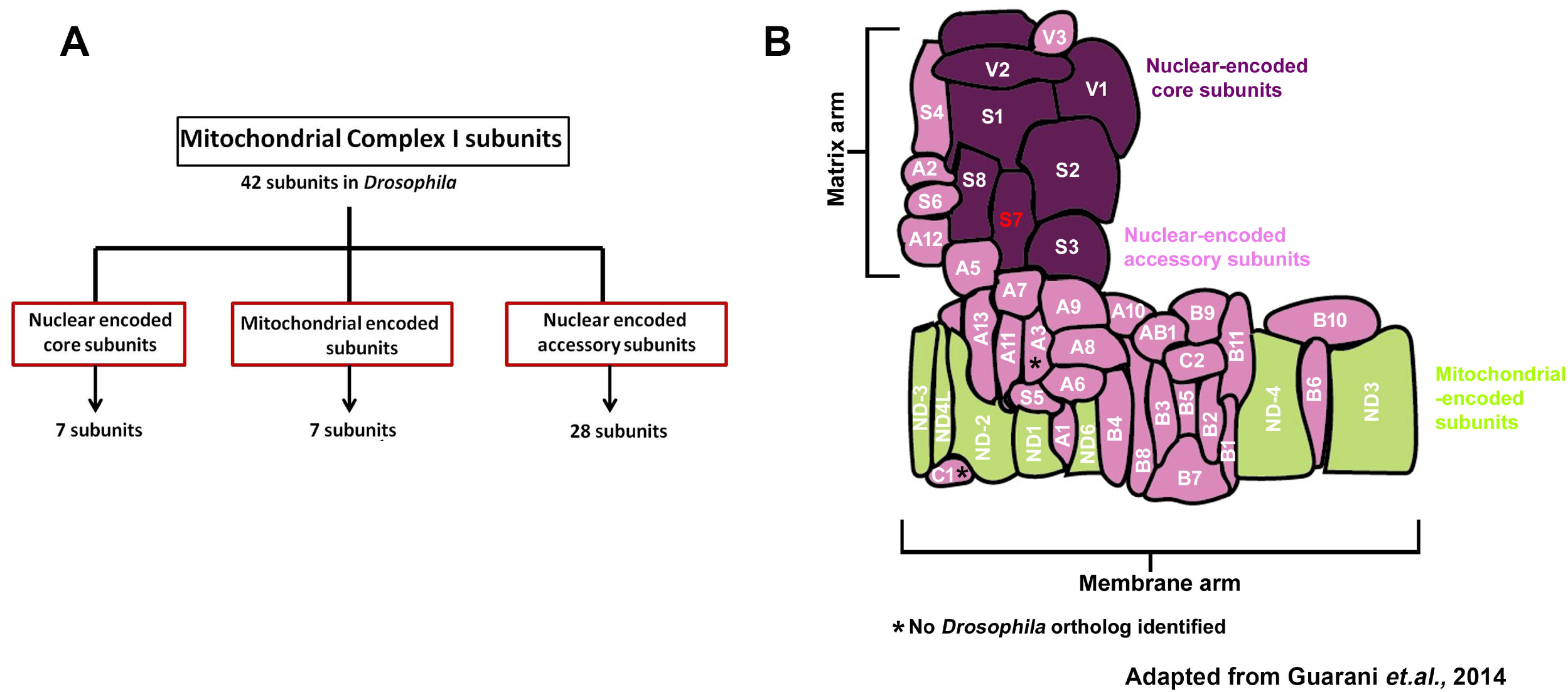
Organization and classification of *Drosophila* Mitochondrial Complex I (Cl) subunits. (A) Schematic showing how Mitochondrial Complex I (MCI) is organized into three distinct classes of subunits (Nuclear-encoded core: 7 subunits, mitochondrial-encoded: 7 subunits and nuclear-encoded accessory: 28 subunits). (B) Schematic representation of how the 42 different subunits of *Drosophila* MCI are arranged to produce the L-shaped topology; adapted from (Guarani et al., 2014). NDUFS7/S7 (*ND-20L*) is labeled in red.

Mitochondrial Complex I subunits have been biochemically characterized from diverse species such as *Bos taurus* (Clason et al., 2010), *Yarrowia lipolytica* (Kashani-Poor et al., 2001; Radermacher et al., 2006), *Pichiapa storis* (Bridges et al., 2009), *Neurospora crassa* (Guenebaut et al., 1997; Leonard et al., 1987), *Drosophila melanogaster* (Garcia *et al*., 2017), humans (Murray et al., 2003) and rodents (Schilling et al., 2005). Electron microscopy experiments revealed that MCI has an L-shaped architecture, including arms that extend into the membrane and the periphery (Clason *et al*., 2010; Guenebaut et al., 1998; Leonard *et al*., 1987; Radermacher *et al*., 2006). Subsequent X-ray crystallography analyses were performed detailing the entire mitochondrial complex from the *Y. lipolytica* at 6.3 Å resolution (Hunte et al., 2010).

Pharmacology in animal models has helped to elucidate some information about MCI activity and physiological consequences of its dysfunction. Two pharmacological agents that target MCI are rotenone and paraquat, which block the flow of electrons from NADH to ubiquinone and trigger ROS formation in cells (Cocheme and Murphy, 2008; Degli Esposti, 1998; Fato et al., 2009). Studies in the fruit fly *Drosophila melanogaster* assessed the effects of rotenone and paraquat on MCI *in vivo* physiology. In flies, paraquat impairs neuronal and mitochondrial function (Hosamani and Muralidhara, 2013). Similarly, delivering rotenone to flies triggers superoxide formation, leading to defects in locomotor ability (Leite et al., 2018). Antidotal pharmacology in *Drosophila* can abate these phenotypes. For example, eicosapentaenoic (EPA) and docosahexaenoic (DHA) omega-3 fatty acids reversed neurotoxic effects in flies induced by paraquat (de Oliveira Souza et al., 2019). Additionally, the sesquiterpene alcohol (-)-α-bisabolol (BISA) successfully reversed rotenone-induced locomotion and lethality phenotypes in flies (Leite *et al*., 2018).

Genetic studies of MCI dysfunction in animals have shed light on possible physiological consequences of mutation of specific subunits (Mayr et al., 2015). One of these subunits, human NDUFS4, is required for MCI assembly, and misregulation of NDUFS4 has been linked with Leigh syndrome and cardiomyopathy (Fassone and Rahman, 2012). Modeling this deficiency in fruit flies, NDUFS4 depletion (*Drosophila* ND-18) leads to a state of progressive neurodegeneration, locomotor defects, and a shortened life span (Foriel et al., 2018). Additionally, among the mitochondrially-encoded MCI subunits, *Drosophila* ND2 loss-of-function mutants display behaviors reminiscent of human mitochondrial disease, including reduced life span and neurodegeneration (Burman et al., 2014). Recently, a genetic study disrupting mouse MCI specifically in dopaminergic neurons showed that loss of MCI alone was sufficient to induce progressive phenotypes reminiscent of Parkinson’s Disease (Gonzalez-Rodriguez et al., 2021).

Despite the existence of these animal models – and despite the association of MCI dysfunction with human neurophysiological and neuromuscular disorders – extensive systematic functional analyses have not been performed to understand MCI roles on a subcellular level in neurons or synapses. Here, we report a small□scale electrophysiology- and RNA interference-based screen examining human neurological and muscle-related genes in *Drosophila* (300 lines). Our screen identified an MCI subunit line whose knockdown in the muscle and neuron led to aberrant neurotransmission. Follow-up electrophysiology showed that genetic depletion of other MCI subunit genes and pharmacology phenocopied this neurotransmission phenotype. On a synapse level, we found that depletion of MCI subunits affected NMJ structure. Together, our data support a role for MCI subunits in synaptic transmission and plasticity. Combined with prior studies, we argue that genetic modeling in fruit flies could be a useful way to understand the mechanisms of MCI subunits in synapse regulation. In turn, this could establish *Drosophila melanogaster* as an organism to further model MCI deficiency diseases.

## 2. Materials and methods

### *Drosophila* husbandry

*Drosophila melanogaster* was cultured on a traditional cornmeal media containing molasses according to a recipe from the Bloomington *Drosophila* Stock Center (BDSC, Bloomington, IN). *Drosophila* husbandry was performed according to standard practices (James et al., 2019). For experiments, larvae were raised at 18°C, 25°C, or 29°C in humidity-controlled and light-controlled (12 hours dark: 12 hours light) Percival DR-36VL incubators (Geneva Scientific).

### *Drosophila* stocks

*w^1118^* (Hazelrigg et al., 1984) was used as a non-transgenic wild-type stock. The *UAS-GluRIII[RNAi]* line was utilized to screen homeostatic candidate molecules as described previously (Brusich et al., 2015; Spring et al., 2016). The Gal4 drivers used simultaneously for the Pre*+*Post-Gal4 conditions were *elaV(C155)-Gal4* (Lin and Goodman, 1994), *Sca-Gal4* (Budnik et al., 1996), and *BG57-Gal4* (Budnik et al., 1996). Many *UAS-RNAi* or genetic mutant lines were obtained from the Bloomington Drosophila Stock Center (Supplementary Tables S1, S2, S3, and S5).

### Larval crawling assay

Vials containing third instar larvae were used for the crawling assay. Vials were supplemented with 4 ml of 20% sucrose solution and left for 10 min to let the larvae float on top. Floating third instar animals were poured into a petri dish and washed gently twice with deionized water in a paintbrush. A minimum 10 larvae of each genotype were analyzed on a 2% agarose gel in a petri dish with gridline markings 1 cm on a graph paper. The larvae were acclimatized in the petri dish before videotaping. The average distance crawled (in centimeters) by larvae was calculated based on the average number of gridlines passed in 30 sec. (Raut et al., 2017).

### Immunohistochemistry

Wandering third instar larvae were dissected and fixed on a sylgard Petri plate in ice-cold Ca^2+^ HL-3 and fixed in 4% paraformaldehyde in PBS for 30 minutes or in Bouin’s fixative for 2 minutes as described earlier (Raut *et al*., 2017). The larvae were washed with PBS containing 0.2% Triton X-100 (PBST) for 30 min, blocked for an hour with 5% normal goat serum in PBST, and incubated overnight in primary antibodies at 4°C followed by washes and incubation in secondary antibodies. Monoclonal antibodies such as anti-DLG (4F3), anti-Synapsin (3C11) were obtained from the Developmental Studies Hybridoma Bank (University of Iowa, USA). All were used at 1:30 dilution. Fluorophore-coupled secondary antibodies Alexa Fluor 488, Alexa Fluor 568 or Alexa Fluor 647 (Molecular Probes, Thermo Fisher Scientific) were used at 1:400 dilution. Alexa 488 or 647 and Rhodamine-conjugated anti-HRP were used at 1:800 and 1:600 dilutions, respectively (Jackson ImmunoResearch Laboratories, Inc.). The larval preparations were mounted in VECTASHIELD (Vector Laboratories, USA) and imaged with a laser scanning confocal microscope (LSM 700; Carl Zeiss). All the images were processed with Adobe Photoshop 7.0 (Adobe Systems, San Jose, CA).

### Confocal imaging, quantification, and morphometric analysis

Samples were imaged using a 700 Carl Zeiss scanning confocal microscope equipped with 63×/1.4 NA oil immersion objective using separate channels with four laser lines (405, 488, 555, and 639 nm) at room temperature. The boutons were counted using anti-Synapsin, anti-HRP co-stained with anti-DLG on muscle 6/7 of A2 hemisegment, considering each Synapsin or HRP punctum to be a bouton. At least 6-8 NMJs of muscle 6/7 (A2 hemisegment) from 4 animals were used for bouton number quantification. All genotypes were immunostained in the same tube with identical reagents for fluorescence quantifications and then mounted and imaged in the same session as we have done previously (Spring *et al*., 2016; Yeates et al., 2017). Z-stacks were obtained using similar settings for all genotypes with z-axis spacing between 0.5-0.7 μm and optimized for detection without saturation of the signal. The Image J software (National Institutes of Health) analysis toolkit used maximum intensity projections for quantitative image analysis. Boutons from muscle 6/7 of A2 hemisegment from at least six NMJ synapses were used to quantify Image J software.

### Relative DLG area quantification

The boutons were stained using anti-HRP and anti-DLG on muscle 6/7 of A2 hemisegment. The total DLG and HRP area were calculated separately using Image J software. A minimum of 35 type-1b boutons from 4 animals (8 NMJs) were used for quantification. The relative DLG area in a bouton is quantified by subtracting the total DLG area with respect to HRP area (Relative DLG area= Total DLG area - Total HRP area).

### Statistical Analyses

A Student’s t-test for pairwise comparisons or a one-way ANOVA with a Tukey’s post-hoc test for multiple comparisons was used for statistical analysis (GraphPad Prism Software). Specific statistical tests are noted in the figure legends and supplementary table files and shown in graphs. The data are presented as mean±s.e.m. The *p*-value of each analysis is indicated in the figures and the supplemental tables. For our electrophysiology screen, our cutoff for a possible hit was two standard deviations evoked amplitude below the control line data, similar to prior electrophysiology screens that we and others have conducted (Brusich *et al*., 2015; Dickman and Davis, 2009; Yeates and Frank, 2021).

### Electrophysiology

All dissections and recordings were performed in a modified HL3 saline (Stewart et al., 1994) containing 70 mM NaCl, 5 mM KCl, 10 mM MgCl2, 10 mM NaHCO3, 115 mM sucrose, 4.2 mM trehalose, 5 mM HEPES, and 0.5 mM CaCl2 (unless otherwise noted), pH 7.2. Neuromuscular junction sharp electrode (electrode resistance between 20-30 MΩ) recordings were performed on muscles 6/7 of abdominal segments A2 and A3 in wandering third-instar larvae as described (James *et al*., 2019). Larval NMJs recordings were performed on a Leica microscope in an HL3 buffer containing 10 mM Mg^2+^ and 0.5 mM Ca^2+^ concentrations using a 10x objective and acquired using an Axoclamp 900A amplifier, Digidata 1440A acquisition system, and pClamp 10.7 software (Molecular Devices). Electrophysiological sweeps were digitized at 10 kHz and filtered at 1 kHz. Data were analyzed using Clampfit (Molecular devices) and MiniAnalysis (Synaptosoft) software. Miniature excitatory post-synaptic potentials (mEPSPs) were recorded without any stimulation and motor axons were stimulated to elicit excitatory post-synaptic potentials (EPSPs). Average mEPSP, EPSP were determined for each muscle and quantal content was calculated by dividing average EPSP amplitude by average mEPSP amplitude. Muscle input resistance (R_in_) and resting membrane potential (V_rest_) were monitored during each experiment. Recordings were rejected if the V_rest_ was above −60 mV and R_in_ was less than 5 MΩ.

### Pharmacology and rotenone feeding assay

For each treatment, either early or late first instar larvae were fed with rotenone dissolved in DMSO (or DMSO alone for controls), allowed to develop into the third instar stage before electrophysiological recordings. Various doses of rotenone ranging from 2 μM to 500 μM were fed to the larvae and sharp electrode recordings were performed at different time points as mentioned in the Figure 4. Additionally, 500 μM of rotenone was acutely applied in an open preparation for 30 minutes before recordings.

## 3. Results

### An electrophysiology-based screen identifies Mitochondrial Complex I subunits in *Drosophila*

RNA interference (RNAi) is a powerful genetic approach. In *Drosophila*, RNAi screens have identified novel genes involved in diverse processes, such as nervous system development, eye development, and wound closure (Koizumi et al., 2007; Lesch et al., 2010; Pignoni et al., 1997; Raut *et al*., 2017; Yamamoto et al., 2014). Prior reports indicate that many human diseases that affect muscle and nervous system health could have primary defects in synaptic function or plasticity (Hirth, 2010). However, the specific functional requirements remain unclear.

Our goal for this study was to survey homologs of selected human disease genes to test if they may also have conserved roles in synapse function or plasticity. We performed a targeted RNAi-mediated reverse genetic screen in *Drosophila*. We picked 300 publicly available *UAS*-driven RNAi lines linked to human neurological and muscle-related disorders, and we knocked down the chosen target genes pan-neuronally and in muscles (Fig. 2A). Because RNAi-mediated knockdown can cause off-target effects or only partial depletion of gene function (Dietzl et al., 2007), we used multiple RNAi lines for genes, whenever possible.

**Figure 2.**
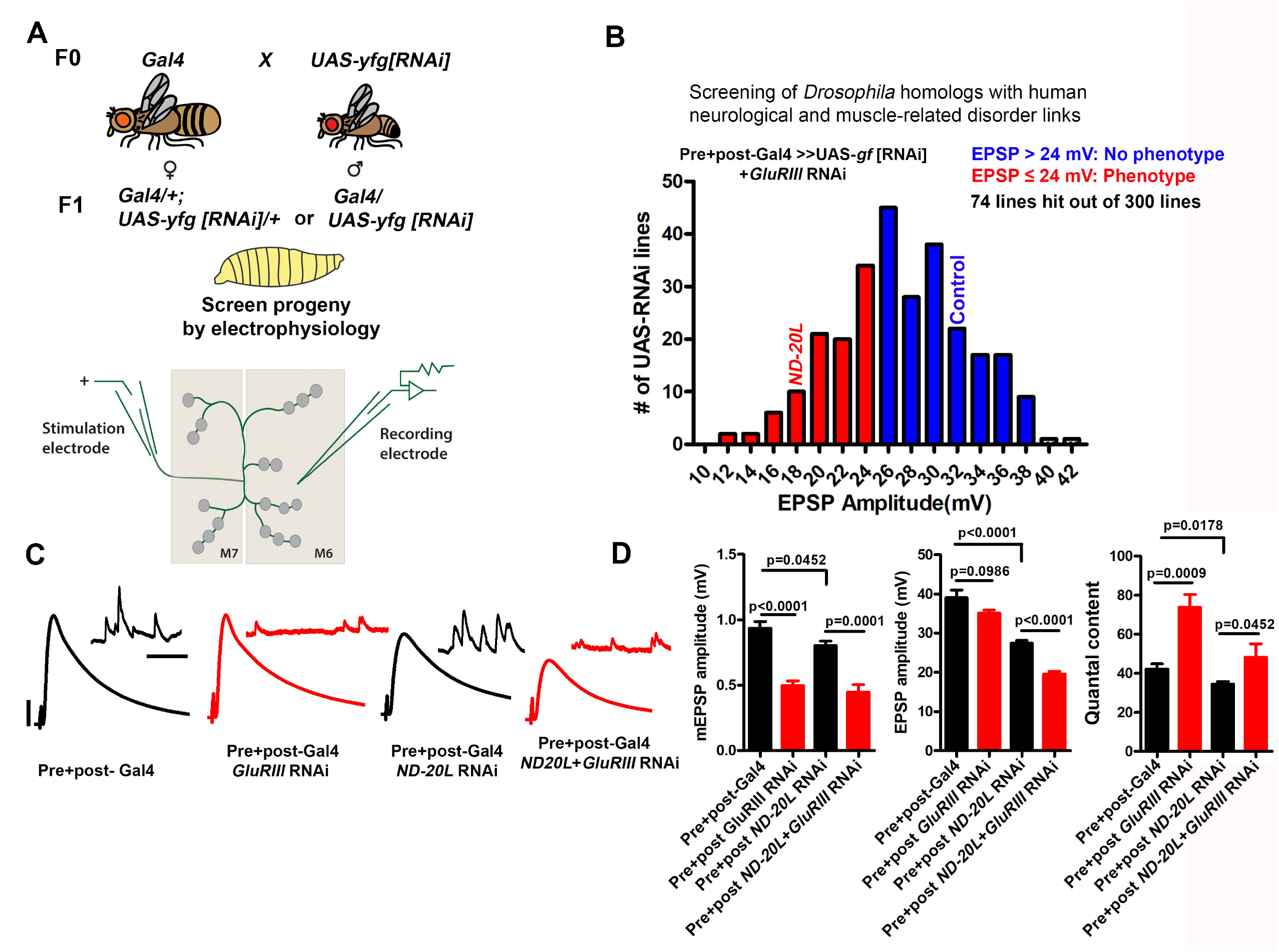
An RNAi screen to identify genes involved in homeostatic synaptic plasticity for human neurological and muscle-related disorders in *Drosophila*. (A) Crossing scheme for screen. Presynaptic + Postsynaptic-Gal4; *GluRIII*[RNAi] × *UAS-yfg* [RNAi] (*yfg*, “your favorite gene”). For UAS-RNAi lines on chromosomes II or III, male progeny were examined by electrophysiology because dosage compensated *elaV(C155)-Gal4/Y* male progeny should have a higher dose of pre-synaptic Gal4 than *elaV(C155)-Gal4/+* female siblings. (B) The mean value of EPSP amplitudes for screened lines. The mean for the control cross x *GluRIII[RNAi]* = 35.22 mV (Table S1, final row), and standard deviation for this control = 5.88 mV. For our analysis, we have included everything as a potential hit which fell roughly two standard deviations below the control mean amplitude or lower (23.46 mV, ~24 mV or lower) (Table S1). Screen results are highlighted with red and blue bars in the histogram. EPSP cutoffs of ≤ 24 mV (red) were considered putative hits, and > 24 mV (blue) were considered normal (Table S1). The majority of knockdown experiments were performed at 29°C, except a few experiments (18°C because higher temperatures caused early lethality). Electrophysiology recordings were carried out at 25°C. (C) Representative electrophysiological traces for selected *ND-20L* or control genotypes in the presence and absence of *GluRIII* RNAi. Scale bars for EPSPs (mEPSPs) are x = 50 ms (1000 ms) and y = 10 mV (1 mV). (D) Quantification showing mEPSP, EPSPs amplitude and quantal content in pre+post-Gal4(mEPSP: 0.93 ± 0.05, EPSP: 38.99 ± 2.02, QC: 42.20 ± 2.65), pre+post-*GluRIII* RNAi (mEPSP: 0.49 ± 0.03, EPSP: 35.08 ± 0.82, QC: 73.82 ± 6.52), pre+post *ND-20L[RNAi]* (mEPSP: 0.80 ± 0.03, EPSP: 27.44 ± 0.70, QC: 34.44 ± 1.34) and pre+post *ND-20L+GluRIII* RNAi (mEPSP: 0.44 ± 0.05, EPSP: 19.55 ± 0.68, QC: 48.18 ± 6.95). The data for pre+post *ND-20L[RNAi]* and pre+post *ND-20L+GluRIII* RNAi are also represented in the supplemental Table S1. Statistical analysis based on one-way ANOVA with Tukey’s post-hoc test. Error bars represent mean±s.e.m. *p*-values are indicated in the figure.

Our main data collection assay was *Drosophila* neuromuscular junction (NMJ) electrophysiology. We recorded spontaneous miniature excitatory postsynaptic potentials (mEPSPs) and excitatory postsynaptic potentials (EPSPs). From these data we were able to assess for each gene knockdown: 1) baseline neurophysiology levels for quantal size, evoked potentials, and quantal content (see Methods); and 2) the maintenance of a form of synaptic plasticity, called presynaptic homeostatic potentiation (PHP). We first assessed neurotransmission and PHP via pre- and post-synaptic knockdown of the selected RNAi line plus knockdown of a glutamate receptor subunit gene (*GluRIII* RNAi), as we have shown previously, as a means of homeostatically challenging synapse function through decreased quantal size (Brusich *et al*., 2015; James *et al*., 2019). The full screen dataset is included in the Supplementary material (Table S1).

Screened Gal4 + RNAi lines with average EPSP amplitudes ≤24 mV were classified as putative hits for this first-pass screen (Figure: 2B). Our cutoff of 24 mV was calculated as approximately two standard deviations below the control data for Gal4 drivers x *GluRIII[RNAi]* (Table S1, final row; mean = 35.22 mV; standard deviation for this control = 5.88 mV). The shortlisted RNAi lines were further analyzed in the absence of the *GluRIII* knockdown in order to differentiate between possible PHP defects versus baseline neurotransmission defects.

The data revealed an intriguing phenotype for a line targeting *ND-20L*, an essential component of Mitochondrial Complex I (electron transport chain of mitochondria). Compared to the driver control alone, pre- + postsynaptic *Gal4* drivers + *UAS-ND-20L[RNAi]* caused a significant defect in baseline evoked neurotransmission (Fig. 2B-D), no defect in quantal size (Fig. 2C-D), and a significant defect in calculated quantal content (QC, Fig. 2D). When presented the homeostatic challenge *UAS-GluRIII[RNAi]* genetic background, *ND-20L* knockdown by RNAi did decrease evoked neurotransmission (EPSP amplitude) significantly further than the non-challenged *ND-20L[RNAi]* control (Fig. 2C-D, Table S2). Yet there was a slight homeostatic increase in QC (Fig. 2D). Together, our screen data suggest that Complex I subunits in *Drosophila* likely regulate baseline synapse function but leave PHP at least partially intact. We tested neurotransmission further.

### Mitochondrial Complex I subunits in neurons and muscle regulate synapse function

ND-20L/NDUFS7 is a core MCI subunit. To assess whether synaptic transmission is also altered after perturbation of other core MCI subunits, we recorded baseline NMJ neurotransmission after knocking down the subunits concurrently in neurons and muscle.

We acquired RNAi-based tools for all core subunits. Baseline control electrophysiology of the Gal4 driver lines alone yielded mEPSPs of 0.75 ± 0.03 mV, EPSPs of 37.67 ± 2.19 mV, and quantal content of 49.70 ± 1.71 (Table S2). For each Complex I RNAi line, the amplitudes of spontaneous miniature postsynaptic potentials (mEPSP) were not significantly altered in pre- + postsynaptic-depleted Complex I subunits (Fig. 3A-S, Table S2). Nor did we find any notable change in mini frequency in pre- + postsynaptic knockdown of Complex I subunits (Table S2). However, for every case the amplitude of evoked postsynaptic potentials (EPSP) was numerically reduced after depleting Complex I, and quantal content (QC) was similarly reduced (Fig. 3A-U, Table S2). This numerical reduction in EPSP amplitude was statistically significant in 11/18 genetic lines we tested (Table S2). Of the 7 lines in which the EPSP reduction did not achieve statistical significance, 4 were mitochondrially-encoded (Table S2). This is not necessarily surprising. RNA interference and message degradation machinery should not target mitochondrially-encoded genes.

**Figure 3:**
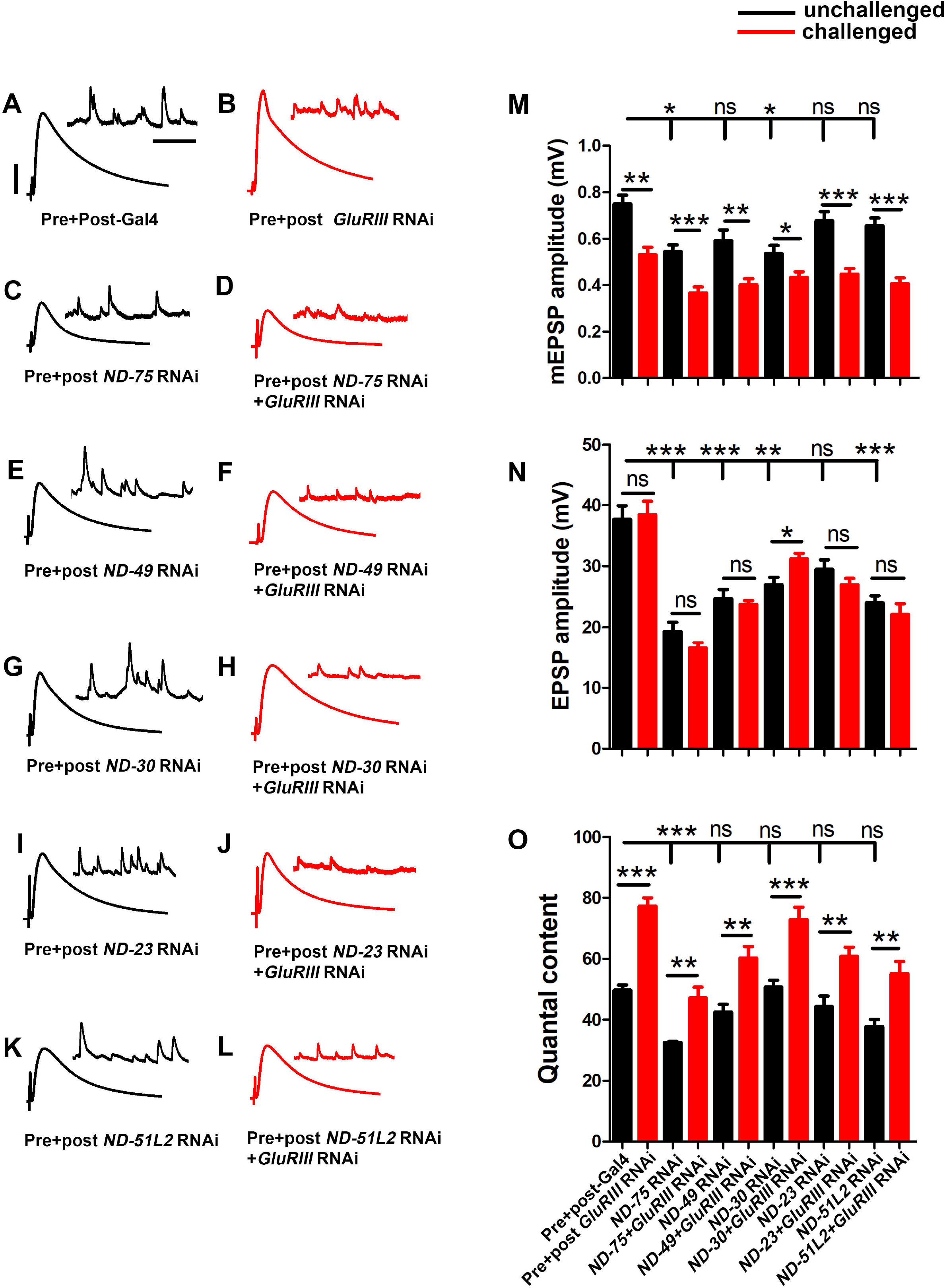
Core complex I subunits regulate neurotransmission. (A-L) Representative traces for pre+post-Gal4 driven knockdown of Mitochondrial Complex I subunits in the presence and absence of *GluRIII* RNAi. Scale bars for EPSPs (mEPSPs) are x = 50 ms (1000 ms) and y = 10 mV (1 mV). (M-O) Histogram showing average mEPSP, EPSPs amplitude and quantal content in pre+post-Gal4 control, pre+post-Gal4 driven *GluRIII* RNAi, pre+post-Gal4 driven MCI subunits and pre+post-Gal4 driven *GluRIII* RNAi and MCI RNAi core subunits in the indicated genotypes. At least 8 NMJ recordings of each genotype were used for quantification. The knockdown experiments and electrophysiology recordings were performed at 25°C. Pre- and postsynaptic RNAi corresponding to MCI subunits were analyzed for EPSPs through electrophysiological recordings. *p*-values for EPSPs are indicated in Table S2. Statistical analysis based on Student’s t-test for pairwise comparisons (in determining homeostatic potentiation for a specific genotype), or one-way ANOVA followed by post-hoc Tukey’s multiple comparisons (if comparing back to control across genotypes in the dataset). Error bars represent mean±s.e.m.

Because of the reduction of neurotransmission in many instances of MCI loss, we further checked if the NMJ could express PHP in the presence of *GluRIII* RNAi. As was the case with *ND-20L[RNAi]*, we found intact PHP signaling in most lines, as indicated by a significant increase in QC after *GluRIII* knockdown (Table S2). Together, our genetic data suggest that a majority of the core Complex I subunits in *Drosophila* regulate baseline synapse function but still have some capacity to express homeostatic plasticity. We decided to characterize MCI function further by tissue specific depletion and pharmacological manipulation.

### Loss of *ND-20L* in muscle affects neurotransmission and larval motor behavior

Next, we depleted MCI using tissue-specific *Gal4* lines. We recorded NMJ electrophysiology for *Gal4* controls, pan neuronal-, or muscle-depleted *ND-20L* by *RNAi* (Fig. 4A-H). For neuronal depletion, the amplitude of spontaneous miniature post-synaptic potentials (mEPSP) was not significantly changed (Fig. 4E), nor were evoked EPSP amplitude or quantal content (Fig. 4G-H).

**Figure 4:**
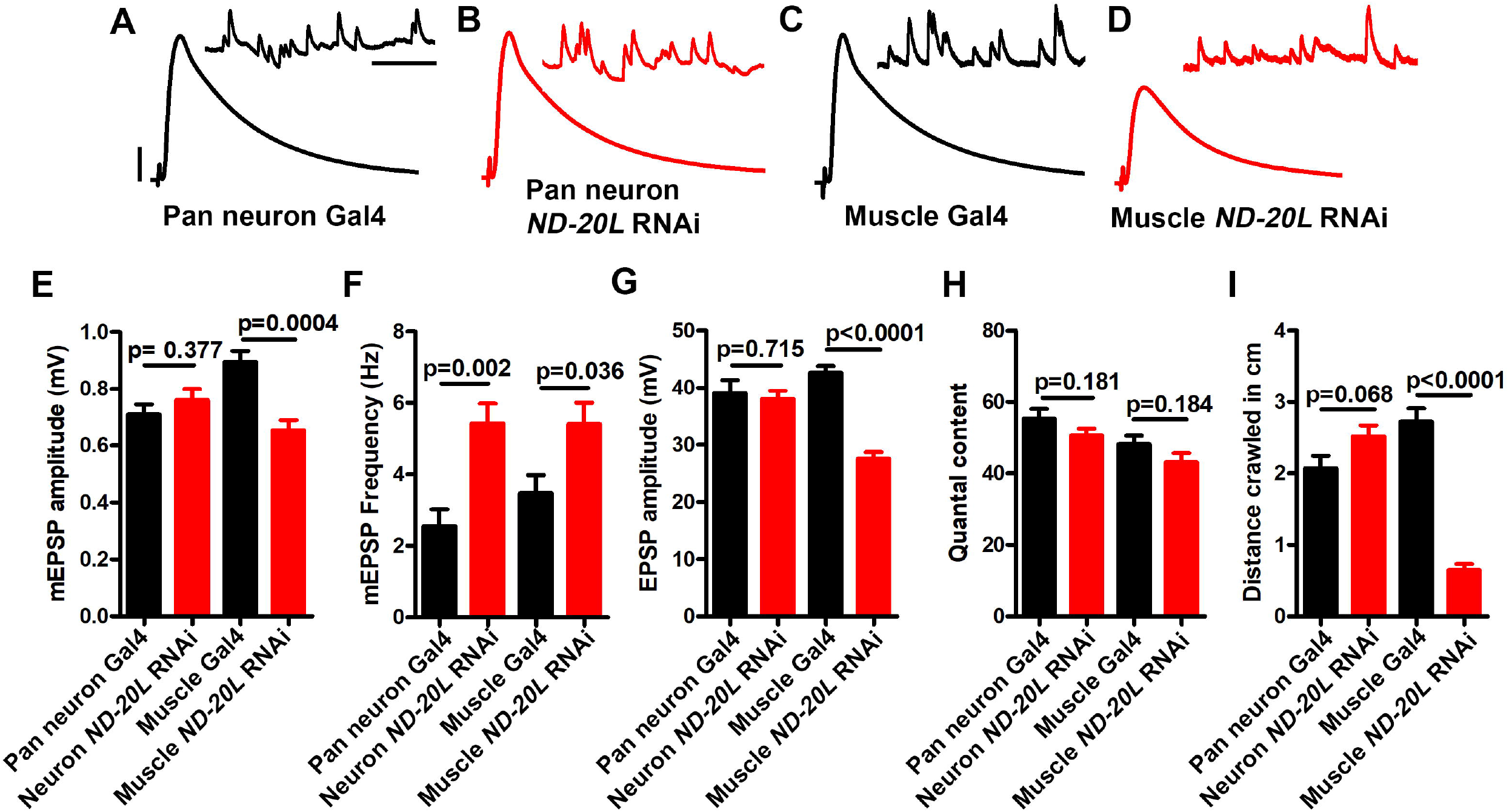
Muscle-specific knockdown of *ND-20L[RNAi]* affects neurotransmission and motor behavior in *Drosophila*. (A-D) Representative traces of mEPSPs and EPSPs in (A-B) pan-neuronal *elaV(C155)-Gal4/+*, pan-neuronal *elaV(C155)^-^Gal4* driven *ND-20L[RNAi] (elaV(C155)/+; ND-20L[RNAi]/+*) and (C-D) *BG-57-Gal4/+*, BG-57-Gal4 driven *ND-20L[RNAi] (BG-57-Gal4/+; ND-20L[RNAi]/+*) animals. Scale bars for EPSPs (mEPSPs) are x = 50 ms (1000 ms) and y = 10 mV (1 mV). The knockdown experiments and electrophysiology recordings were performed at 25°C. Note that EPSPs amplitudes were reduced in muscle depleted *ND-20L[RNAi]* while pan-neuronal depleted *ND-20L* larvae did not show any remarkable change in EPSPs. (G-J) Histograms showing average mEPSPs amplitude, mEPSP frequency, EPSPs amplitude and quantal content are indicated in the table. At least 6 NMJs recordings of each genotype were used for quantification. (I) Histogram showing crawling behavior (in cm) of the larvae in the indicated genotypes. Knocking down *ND-20L[RNAi]* in muscle showed a severe defect in crawling behavior. However, neuronally depleting *ND-20L* did not show any significant crawling defects. At least 10 animals were analyzed for crawling behavioral analysis. p-values are indicated in the figure. Statistical analysis based on Student’s t-test for pairwise comparison. Error bars represent mean±s.e.m.

By contrast, the mEPSP amplitude and the EPSP amplitude were both significantly decreased in the muscle-depleted *ND-20L[RNAi]* NMJs (Fig. 4 C-H) (Raw data, Table S3). This concurrent decrease in both mEPSP and EPSP amplitudes yielded a calculated quantal content that was not significantly different for the muscle-depleted *ND-20L[RNAi]* NMJs (Fig. 4H). Interestingly, there was an increase in mEPSP frequency in for both tissue-specific depletion conditions (Fig. 4F).

Additionally, we performed behavioral experiments on pan neuronal- and muscle-depleted *ND-20L[RNAi]* third instar larvae. Consistent with the electrophysiological recordings, muscle-depleted animals showed severe defects in crawling ability while pan-neuronal depletion of *ND-20L* did not show any significant defects in crawling behavior (Fig. 4I, Table S3).

### Pharmacology phenocopies Mitochondrial Complex I subunit gene knockdown

Prior studies in *Drosophila* have targeted MCI via pharmacology. This has resulted in neurological and behavioral defects in adult flies reminiscent of human neurodegenerative conditions (Leite *et al*., 2018). If it is the case that our genetic knockdown experiments are specific for MCI, then we should be able to phenocopy the NMJ electrophysiological defects with pharmacology. To target MCI, we used rotenone. Rotenone is an isoflavone derived from plants, and it directly disrupts the electron transport chain of MCI and can result in an accumulation of reactive oxygen species (ROS) (Degli Esposti, 1998; Fato *et al*., 2009).

We used several methods to deliver rotenone to the NMJ. We varied multiple parameters: drug concentration, exposure time, and delivery method. Larvae fed a low dose of rotenone (2 μM) for 48 hours showed no defect in evoked EPSP amplitude compared to carrier alone (5% DMSO, 48 hours, Figs. 5A-B, N). Likewise, freshly dissected NMJ preps acutely soaked in a high dose of rotenone (500 μM) showed little EPSP defect compared to carrier alone (5% DMSO, Figs. 5C-D, N).

**Figure 5:**
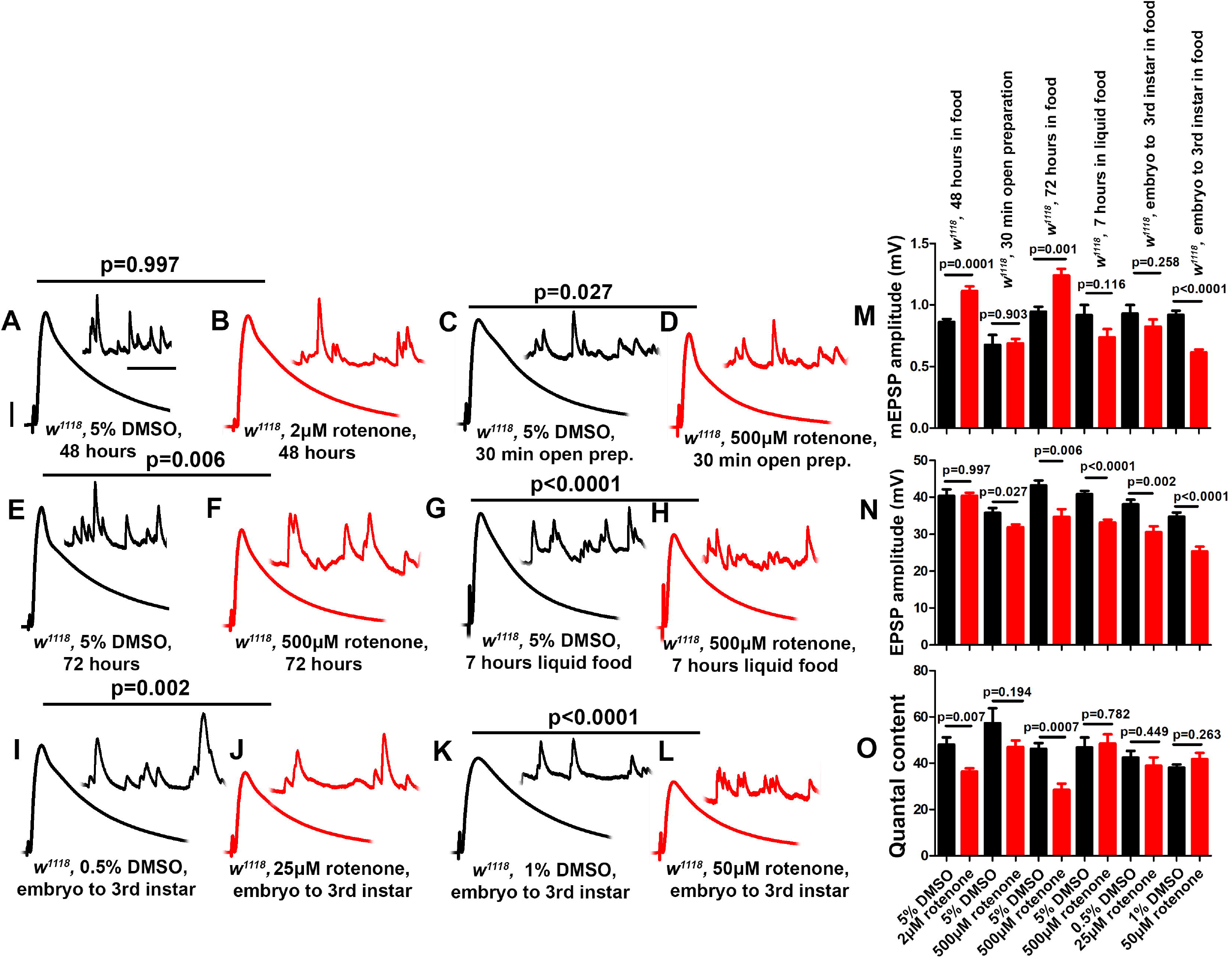
Rotenone diminishes neurotransmission at the NMJ. (A-L) Representative traces for third instar wild-type larvae treated or fed with DMSO and rotenone at various concentrations as indicated above. Scale bars for EPSPs (mEPSPs) are x = 50 ms (1000 ms) and y = 10 mV (1 mV). (M-O) Histogram showing average mEPSP, EPSPs amplitude and quantal content of larvae treated with DMSO and rotenone in the indicated conditions. Note that larvae treated or fed a high dose of rotenone showed decreased EPSP amplitudes compared to carrier controls. The flies were grown in a molasses media and recordings were performed at 25°C. At least 8 NMJs recordings of each genotype were used for quantification. *p*-values for EPSPs are indicated in Table S4. Statistical analysis based on Student’s t-test for pairwise comparison. Error bars represent mean±s.e.m.

However, larvae fed a high dose of rotenone (500 μM) for longer periods (72 hours and 7 hours) showed blunted EPSP amplitudes compared to carrier controls (Fig. 5E-H, N). Additionally, larvae fed intermediate doses (25 μM or 50 μM) throughout life (approximately 120 hours from egg laying to third instar) also showed blunted EPSP amplitudes (Fig. 5I-L, N).

Interestingly, not all of the delivery methods that caused a diminishment of EPSP amplitudes resulted in a decrease in quantal content (Fig. 5M, O). This is because some delivery methods concurrently changed both quantal amplitude (Fig. 5M) and evoked amplitude (Fig. 5N). This was similar to what we observed for postsynaptic knockdown of *ND-20L* (Fig. 4). One possible reason is that application of a drug that impairs MCI could also have postsynaptic effects that govern sensitivity to individual vesicles of neurotransmitter.

### Mitochondrial Complex I subunits regulate NMJ structural plasticity

We conducted NMJ staining experiments to assess if cytological phenotypes might accompany MCI loss. While Complex I subunits are expressed ubiquitously throughout the development, their roles in synapse development are largely unknown. To test this parameter, we knocked down some of the Complex I core subunits that we had previously analyzed electrophysiologically (using pre+post synaptic Gal4 drivers), but this time we analyzed the larval neuromuscular junction (NMJ) morphology. To visualize NMJ bouton morphology, we co-stained with anti-Synapsin (a pre-synaptic marker) and anti-Discs Large (DLG, a post-synaptic marker) (Benson and Voigt, 1995; Budnik *et al*., 1996).

Pre- + postsynaptic knockdown of Complex I subunits caused diminished NMJ structural development (Fig. 6). For each subunit targeted, we observed diminished NMJ growth (Fig. 6A-F). This phenotype manifested in two different ways for different subunits: either as a decrease in total bouton number for animals reared at 25°C (Fig. 6F) or as a diminishment in bouton area (*ND-20L[RNAi]*) (Fig. 6G-I).

**Figure 6:**
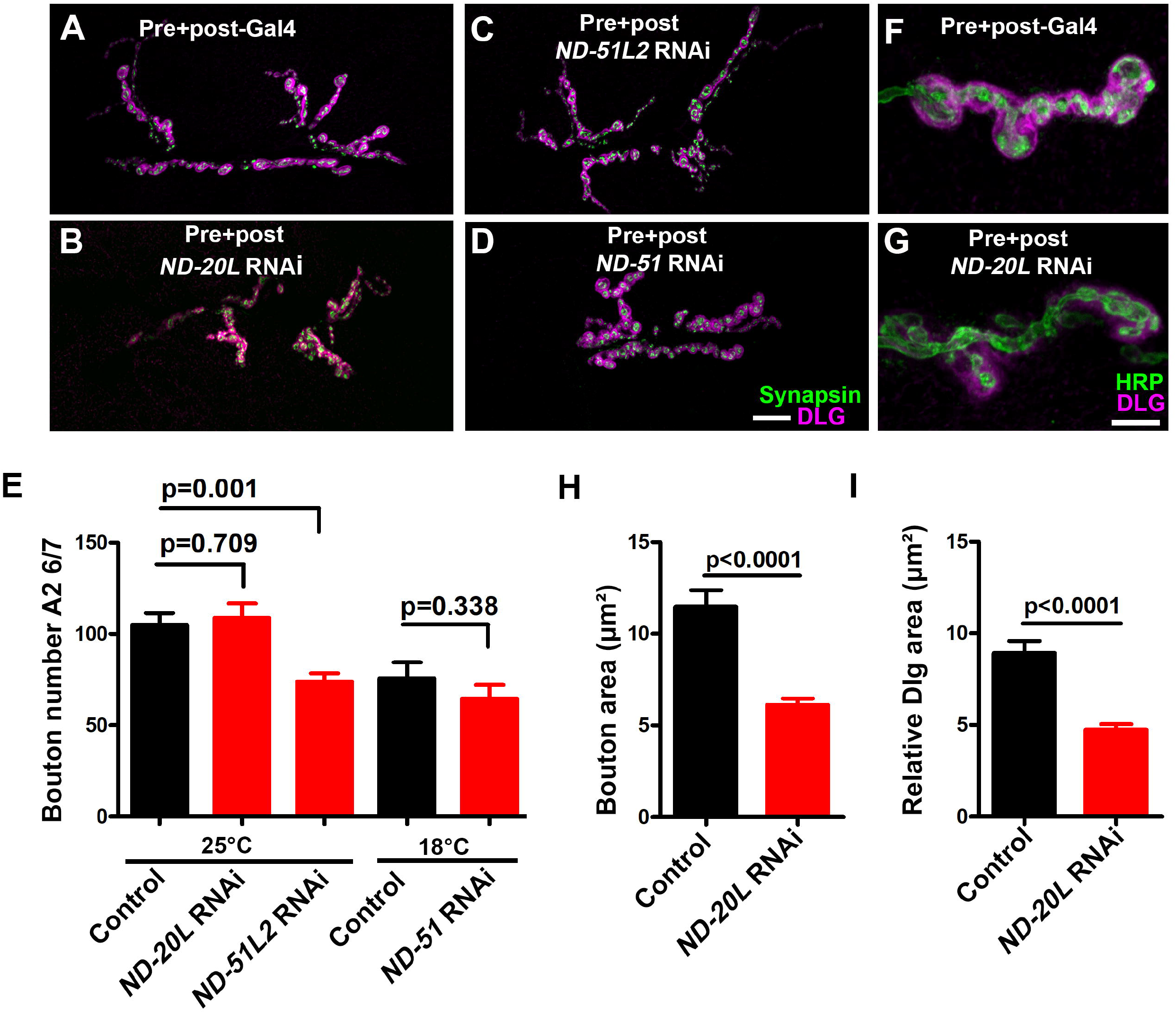
*Drosophila* Mitochondrial Complex I subunits are required for normal neuromuscular junction (NMJ) morphology. (A-D) Representative confocal images of NMJ synapses at muscle 6/7 of (A) Pre+post-Gal4 control (105.0 ± 6.38), Pre+post-Gal4 driven (B) *ND-20L[RNAi]* (108.9 ± 8.01), (C) *ND-51L2* RNAi (73.88 ± 4.50) and (D) *ND-51* RNAi (64.63 ± 7.46) flies double immunolabeled with DLG (magenta) and synapsin (green) antibodies. The NMJ morphological defects were observed in the indicated genotypes. The flies were reared at 25°C. The knockdown of *ND-51* RNAi was lethal in the third instar larval stage at 25°C and 29 °C. Hence we have conducted this experiment at 18°C with respective control. Scale bar: 10 μm. (E) Histograms showing the average total number of boutons at muscle 6/7 of A2 hemisegment in Pre+post-Gal4 control (105.0 ± 6.38), Pre+post-Gal4 driven *ND-20L[RNAi]* (108.9 ± 8.01), *ND-51L2* RNAi (73.88 ± 4.50), control (75.86 ± 8.57) and *ND-51* RNAi (64.63 ± 7.46) larvae. (F-G) Confocal images of boutons at third instar larval NMJ synapse in (F) Pre+post-Gal4 control and (G) Pre+post-Gal4 driven *ND-20L[RNAi]* double immunolabeled with DLG (magenta) and HRP (green) antibodies. Note that the gross morphology of SSR and the immunoreactivity of DLG were reduced in *ND-20L[RNAi]* compared to control. (I-J) Histograms showing average bouton area (I) relative DLG area (J) in μm of Pre+post-Gal4 control (bouton area: 11.49 ± 0.88, relative DLG area: 8.93 ± 0.65), Pre+post-Gal4 driven *ND-20L[RNAi]* bouton area: (6.12 ± 0.34, relative DLG area: 4.74 ± 0.29) larvae. Scale bar represents 10 μm. At least 35 boutons of muscle 6/7 of A2 hemisegment from 4 animals (8 NMJs) were used for area quantification. Statistical analysis based on Student’s t-test for pairwise comparison and one-way ANOVA with Tukey’s post-hoc test for multiple comparisons. Error bars represent mean±s.e.m. *p*-values are indicated in the figure.

For the case of *ND-20L[RNAi]* the decreased bouton size was coupled with a collapsing postsynaptic density, as indicated by decreased amounts of DLG (Fig. 6G-J). Additionally, pharmacological blockade of MCI activity showed similar phenotypes to those of genetic knockdown (Fig. 7 A-J). Together, these data indicate that Complex I subunits regulate NMJ growth and morphology in *Drosophila*.

**Figure 7:**
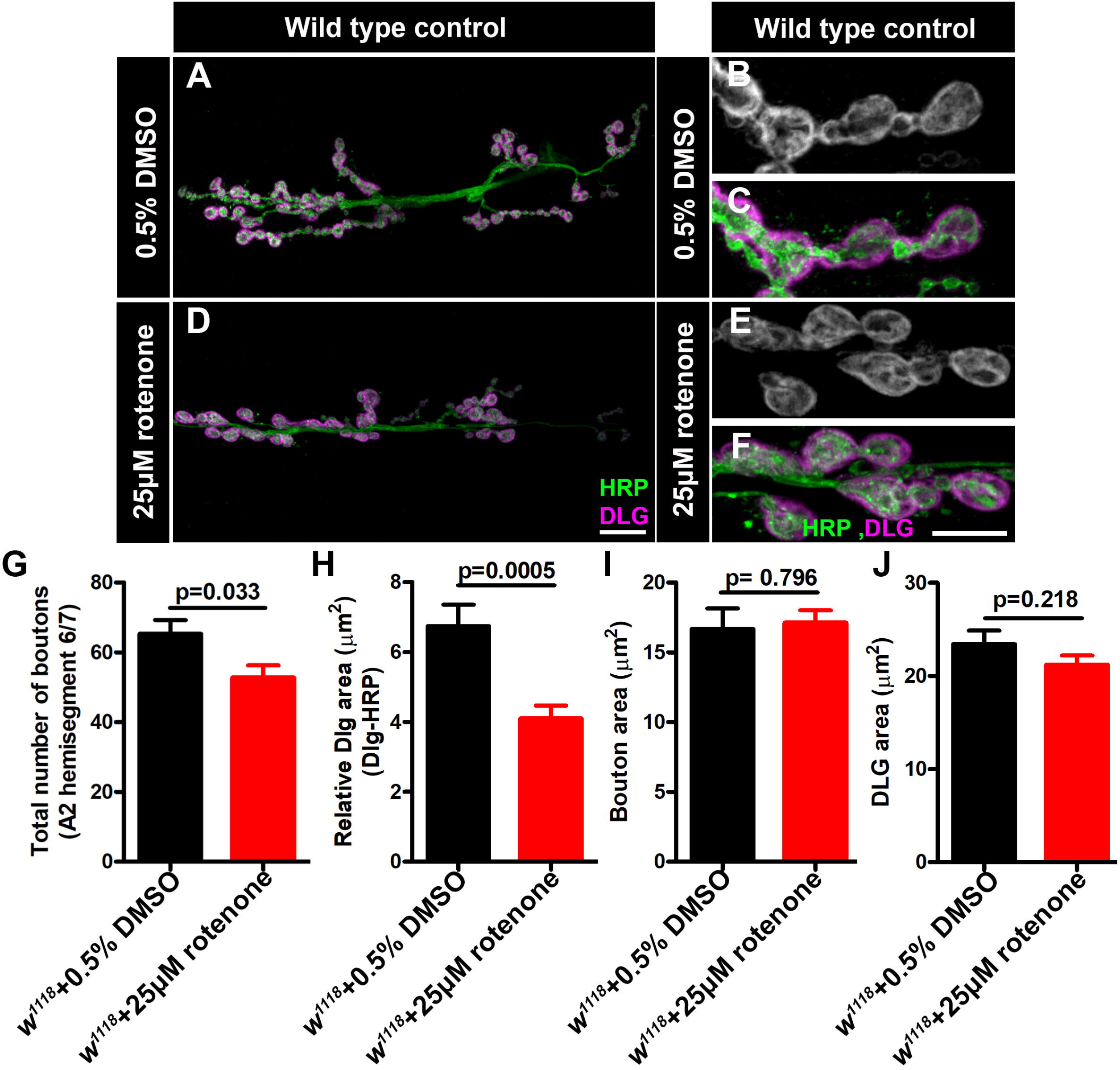
Rotenone affects neuromuscular junction (NMJ) morphology. (A-F) Representative confocal images of NMJ synapses at muscle 6/7 of (A-C) DMSO control, (D-F) Larvae fed with 25 μM rotenone double immunolabeled with DLG (magenta) and HRP (green) antibodies. The NMJ morphological defects were observed in the indicated genotypes. The flies were reared at 25°C to conduct NMJ analysis. Scale bar: 10 μm. (G-J) Histograms showing the average total number of boutons, relative DLG area, bouton area and DLG area at muscle 6/7 of A2 hemisegment in (G) DMSO control (65.25 ± 3.976, N=8), Larvae fed with 25μM rotenone (52.75 ± 3.514, N=8), (H) DMSO control (6.742 ± 0.6156, N=30), Larvae fed with 25μM rotenone (4.101 ± 0.3706, N=30), (I) DMSO control (16.67 ± 1.461, N=30), Larvae fed with 25μM rotenone (17.11 ± 0.8969, N=30) and (K) DMSO control (23.41 ± 1.473, N=30), Larvae fed with 25μM rotenone (21.21 ± 0.9790, N=30) samples. Rotenone-treated larvae showed reduced boutons number and DLG area at the NMJs. Minimum 35 type-1b boutons from 4 animals (8 muscle 6/7 NMJs of A2 hemisegment) were used for quantification. p-values are indicated in the figure. Statistical analysis based on Student’s t-test for pairwise comparison. Error bars represent mean±s.e.m.

## 4. Discussion

*Drosophila* Mitochondrial Complex I subunits are highly conserved with their mammalian counterparts, potentially making *Drosophila* an ideal system to understand cellular MCI roles in diseases (Fig. 1). In this study, we provide a snapshot of synaptic phenotypes that result from MCI impairment in *Drosophila*. These initial studies should help us to dissect the function of MCI in nervous system physiology. Our aggregate data are consistent with the idea that normal MCI enzyme function is required to support normal levels of synaptic transmission (Figs. 2-5). Diminished synaptic neurotransmission may be related to blunted synapse development (Fig. 6-7). Notably, our current study also defines some cell-specific roles that MCI is playing in pre- and postsynaptic compartments. In the context of human disease, genetic MCI loss is global and not tissue-specific. For future our studies, the genetic toolkit available for *Drosophila* means that it should be possible to define additional roles of MCI, in presynaptic neurons, postsynaptic muscle, or in nearby tissue like glia.

### Prior studies are consistent with synaptic functions of Complex I

Genetic dysfunction of MCI subunits produces superoxide in the mitochondrial matrix (Antonucci et al., 2019). Superoxide radicals can be converted into hydroxyl radicals that are highly reactive and cause cellular damage. Pharmacological perturbation of Complex I with rotenone or paraquat stimulates ROS production both *in vivo* and *in vitro* (Cocheme and Murphy, 2008; Degli Esposti, 1998; Fato *et al*., 2009). Relatedly, in adult *Drosophila*, Complex I inhibition by rotenone and paraquat triggers an acute response that is reminiscent of human mitochondrial disorder phenotypes (Hosamani and Muralidhara, 2013).

Although several genetic and biochemical studies have been performed to assess the efficacy of various compounds in suppressing rotenone and paraquat-induced toxicity in flies, little progress has been made to understand the precise consequences of Complex I dysfunction in nervous system physiology. Human mitochondrial disease-causing mutations have been identified in 33/44 MCI subunits (Mayr *et al*., 2015). Mutation of *NDUFS4* causes Leigh syndrome and cardiomyopathy (Fassone and Rahman, 2012). In *Drosophila, dNDUFS4* depletion in neurons showed progressive neurodegeneration, shortened life span and locomotory defects, thus recapitulating patients with *NDUFS4* dysfunction (Foriel *et al*., 2018). Additionally, ubiquitous knockdown of Drosophila *dNDUFS7 (ND-20*) and *dNDUFV1 (ND-51*) causes pupal eclosion defects and alternation in life span (Foriel et al., 2019). Many of these phenotypes are consistent with possible synapse dysfunction.

Recently, a *Drosophila* model of Complex I deficiency was created for mitochondrially encoded subunit ND2 (Burman *et al*., 2014). The *dND-2* mutants showed hallmarks of mitochondrial diseases, including progressive neurodegeneration, muscle degeneration and reduced life span (Burman *et al*., 2014). Other reports have highlighted the role of neuron and glial cells in the context of neurodegeneration triggered by Complex I inhibition (Hegde et al., 2014). For instance, simultaneous disruption of *NDUFS1* in neurons and glia showed progressive neurodegeneration in flies, suggesting that both cell populations are essential in *Drosophila* (Hegde *et al*., 2014). By contrast, depletion of *dNDUFS8* in glial cells did not affect longevity and locomotor ability in animals, while exhibiting significant neurodegeneration in the brain (Cabirol-Pol et al., 2018). A very similar phenotype was observed for *dNDUFS7B* where knockdown in neuron causes increased aggression while exhibiting no effect when depleted in glia (Li-Byarlay et al., 2014). Another study reveals that disruption of MCI subunits in flight muscle impairs MCI assembly (Garcia *et al*., 2017). Moreover, *dNDUFS7 (ND-20L* and *ND-20*) belongs to core subunits, essential for the stability and function of MCI. Hence, we speculate that it might affect both the biogenesis and morphology of mitochondria in *Drosophila*.

### Limitations and Considerations

There are some limitations to the present study. First, any screening endeavor is a first-pass inquiry. Based on our levels of analyses, we cannot definitively rule out roles in neurotransmission or homeostatic plasticity for factors that did not meet our threshold for further study (Fig. 2, Table S1). Conversely, we cannot definitively pinpoint why we saw the neurotransmission defects we saw for the majority of factors that did meet our cutoff (Table S1). We are actively following up on some of these for future work.

Second, we were surprised that our screen had such a high “hit” rate. 74/300 lines screened showed some level of neurotransmission defect that fell more than two standard deviations below what was acquired for the control data set. This high rate could be a function of selection bias. We conducted our analyses on *Drosophila* homologs of human genes previously linked to neurological or muscular diseases. Therefore, this group might be over-represented with factors that are linked to synapse function. Additionally, we used *UAS-RNAi* lines as a screening tool. A recent study demonstrated that RNAi lines with an *attP40* insertion site may have a reduction in adjacent *ND-13A* expression, and *ND-13A* encodes an MCI accessory subunit (Groen et al., 2022). For our work, this could mean that any *attP40* lines are sensitized to MCI impairment and potentially lower neurotransmission. It could explain our puzzling finding that some of the mitochondrial-encoded MCI subunit RNAi lines also showed marginally decreased neurotransmission (Table S2). Those *attP40* lines might actually have a concurrent impairment of *ND-13A*, which we identified in our screen as well (Table S1).

Finally, we note variability in some of our electrophysiology or synapse imaging phenotypes based on the MCI subunit targeted. The subunits that were included in our analyses are essential for maintaining the stability and assembly of MCI. The variability in phenotypes among the subunits could be due to different levels of knockdown of the MCI. It could also be due to the fact that factors like ND-20L seem to have physiological roles in the neuron and the muscle, potentially affecting both presynaptic neurotransmission and glutamate receptor function.

### Possible Models: presynaptic release or postsynaptic ROS

Despite prior genetic and biochemical studies in the different model organisms, little is known about the consequences of MCI dysfunction in nervous system at a single synapse level. At this point, we can propose a couple of different possibilities for impaired NMJ function and morphology. By one model, we can consider presynaptic compartment and synaptic vesicle fusion. Synaptic vesicle fusion is an energy-dependent process and depletion of Complex I subunits disrupt the transfer of electrons from NADH to ubiquinone. Hence, decreased proton motive force would reduce the amount of cellular ATP by oxidative phosphorylation available for release. By another model, we speculate that ROS at the nerve terminal could affect the stability and function of microtubules and microtubule-associated proteins, which are essential for synapse function. The fact that our neuron-specific knockdowns of *ND-20L* did not markedly affect neurotransmission (Fig. 4) could mean that knockdown in neurons was not effective enough, or it was compensated by unknown mechanisms.

By another model, disruption of Complex I in the muscle might produce elevated levels of reactive oxygen species intermediate (ROS), which are reactive and oxidizing. We speculate that ROS formation in the postsynaptic compartment might oxidize and disrupt postsynaptic structures such as DLG (Fig. 6-7), Spectrin, and glutamate receptors that are crucial to maintaining a stable synaptic connection. The alteration of NMJ morphology could be a secondary consequence of the loss of postsynaptic structures, but it could also have a profound effect on synapse function. Another possibility is that MCI dysfunction in the muscle could alter mitochondria biogenesis and diminish ATP production. Assuming that muscle ATP is essential for normal maintenance of muscle health, and any alteration of it might have a severe consequence on the organization of the post-synaptic cytoskeleton, glutamate receptor clusters that are critical for stable synapse function. Our near-future studies on Complex I subunits will dissect distinct signaling mechanisms for *Drosophila* synapse regulation and function. In turn, studies downstream can test if findings in our fruit fly model extend to vertebrate synaptic systems or mitochondrial disease models.

## 7. Conflicts of interest

The authors declare that the research was conducted in the absence of any commercial or financial relationships that could be construed as a potential conflict of interest.

## 8. Author contributions

B.M. and C.A.F. designed the research; B.M. performed the research; B.M. and C.A.F. analyzed the data; B.M. and C.A.F. wrote the paper.

## 9. Funding

This work was supported by a grant from the National Institutes of Health/NINDS (R01NS085164) to C.A.F. B.M. was supported in part by this grant.

## 10. Acknowledgments

We acknowledge the Developmental Studies Hybridoma Bank (Iowa, USA) for antibodies used in this study and the Bloomington Drosophila Stock Center for fly stocks. We thank Frank lab members for their helpful comments and discussions in this study. For research in progress discussions, we thank members of the laboratories of Drs. Tina Tootle, Toshihiro Kitamoto, Pamela Geyer, and Lori Wallrath; we also thank faculty from the Department of Anatomy and Cell Biology at the University of Iowa for weekly workshop discussions. An earlier version of this manuscript was released as a pre-print at *bioRxiv*, (Mallik and Frank, 2021).

## 11. Supplementary Material

Please see Supplementary Tables S1-S5. Table S1 details the genetic screen. Tables S2 and S3 show raw electrophysiology data for Mitochondrial Complex I loss-of-function conditions. Table S4 shows raw electrophysiology data for rotenone application. Table S5 details *Drosophila* stocks and antibodies used.

## 12. Data Availability Statement

The raw data supporting the conclusions in this article will be made available by the authors, without undue reservation.

## Notes

### Competing Interest Statement

The authors have declared no competing interest.

### Summary of Updates

This is a revised version of our manuscript after receiving peer review feedback.

