## Supplementary material for "Roles for Mitochondrial Complex I subunits in regulating synaptic transmission and growth": Table S1

| **Sr. No.** | **Gene name** | **Stock number** | **Mean value of**  **EPSPs in mV** | **Stock number** | **Mean value of EPSPs in mV** |
| --- | --- | --- | --- | --- | --- |
| **A** | ***Muscular Dystrophies*** |  |  |  |  |
| **1** | *CG15651* | BL 29616 | 19.35 | BL 51889 | 28.07 |
| **2** | *syntrophin-like 1* | BL 27504 | 30.56 | BL 61987 | 29.23 |
| **3** | *wing blister* | BL 29559 | 19.35 |  |  |
| **4** | *sarcoglycan alpha* | BL 34027 | 24.70 |  |  |
| **5** | *dystroglycan* | BL 34895 | 22.92 |  |  |
| **6** | *twisted* | BL 55735 | 23.07 |  |  |
| **7** | *dystrobrevin* | BL 32935 | 23.02 | BL 36101 | 31.70 |
| **8** | *dlg1* | BL 34854 | 24.02 | BL 36771,35286 | lethal |
| **9** | *bent* | BL 31545 | 21.69 | BL 31546 | 28.13 |
| **10** | *sarcoglycan beta* | BL 29551 | 30.87 |  |  |
| **11** | *syntrophin-like 2* | BL 42601 | 21.60 | BL 28363 | 21.95 |
| **B** | ***Seizure disorders/Epilepsy*** |  |  |  |  |
| **12** | *couch potato* | BL 60388 | 37.50 | BL 28360 | 29.38 |
| **13** | *jitterbug* | BL 39070 | 23.99 | BL 31590 | lethal |
| **14** | *julius seizure* | BL 28764 | 26.15 |  |  |
| **15** | *technical knockout* | BL 38251 | 20.10 |  |  |
| **16** | *kazachoc* | BL 34584 | 24.34 |  |  |
| **17** | *shaker* | BL 53347 | 30.85 | BL 31680 | 32.57 |
| **18** | *para* | BL 31471 | 36.42 | BL 31676 | 27.95 |
| **19** | *Letm1* | BL 37502 | 32.33 |  |  |
| **20** | *slowpoke* | BL 31677 | 17.15 | BL 26247 | 29.94 |
| **21** | *easily shocked* | BL 38528 | 24.51 |  |  |
| **22** | *ether a go-go* | BL 31679 | 26.74 |  |  |
| **23** | *knockdown* | BL 36740 | 26.00 |  |  |
| **24** | *shaking B* | BL 27292 | 23.53 | BL 27291 | 27.82 |
| **25** | *seizure* | BL 31681 | 24.02 | BL 31682 | lethal |
| **C** | ***Mitochondrial complex deficiency diseases*** |  |  |  |  |
| **26** | *ND-13B* | BL 16860 | lethal |  |  |
| **27** | *mt:ND4L* | BL 65158 | 20.09 |  |  |
| **28** | *ND-49* | BL 28573 | 23.71 | BL 57499 | 20.80 |
| **29** | *mt:ND3* | BL 77153 | 24.33 |  |  |
| **30** | *mt:ND5* | BL 77336 | 25.48 |  |  |
| **31** | *ND-20L* | BL 62381 | 19.08 |  |  |
| **32** | *ND-75* | BL 33911 | 16.53 | BL 27739 | 18.25 |
| **33** | *CG3270* | BL 31153 | 34.46 | BL 58091 | 31.43 |
| **34** | *CG3803* | BL 31073 | 28.20 |  |  |
| **35** | *ND-B17.2* | BL 36695 | 19.01 |  |  |
| **36** | *CG11722* | BL 52920 | 30.47 | BL 28567 | 31.25 |
| **37** | *tetratricopeptide repeat domain 19* | BL 64958 | 28.64 |  |  |
| **38** | *sicily* | BL 55332 | 23.86 |  |  |
| **39** | *ND-13A* | BL 28576 | 16.81 | BL 51860 | lethal |
| **40** | *lethal (2)k14505* | BL 53352 | lethal |  |  |
| **41** | *lethal (2)37Bb* | BL 42608 | 25.97 | BL 3579 | 38.26 |
| **42** | *ND-B12* | BL 61321 | 23.69 |  |  |
| **43** | *synthesis of cytochrome c oxidase* | BL 55179 | 25.08 |  |  |
| **44** | *CG8067* | BL 31102 | 27.43 | BL 31240 | 29.22 |
| **45** | *ND-23* | BL 30487,30201 | 21.93, 18.56 | BL 51797 | 26.95 |
| **46** | *lethal (2)37Bb* | BL 42608 | 25.97 | BL 3579 | 38.26 |
| **47** | *CG5037* | BL 31071 | 26.04 | BL 31169 | 25.68 |
| **48** | *bicoid stability factor* | BL 31078 | 22.48 | BL 31097 | 25.93 |
| **49** | *bcs1 chaperone* | BL 31074 | 33.74 | BL 31075 | 27.58 |
| **50** | *surfeit 1* | BL 31210 | 24.62 | BL 31179 | 27.63 |
| **51** | *ND-42* | BL 32998 | 19.47 | BL 34526 | 16.59 |
| **52** | *ND-51L2* | BL 64536 | 22.07 |  |  |
| **53** | *mt:ND-6* | BL 64670 | 25.27 |  |  |
| **54** | *ND-20* | BL 64995 | 27.71 |  |  |
| **55** | *mt:ND-1* | BL 64852 | 30.37 |  |  |
| **56** | *ND-51* | BL 36701, 29534 | 15.82, 24.78 | BL 52939 | 30.18 |
| **57** | *ND-30* | BL 44535 | 31.17 | BL 51425 | 33.94 |
| **58** | *ND51L1* | BL 42591 | 30.88 |  |  |
| **59** | *ND-24* | BL 51855 | 26.38 |  |  |
| **60** | *mt:ND-4* | BL 65216 | 29.91 |  |  |
| **61** | *mt:ND-2* | BL 77154 | 29.75 |  |  |
| **62** | *ND-24L* | BL 60873 | 29.41 |  |  |
| **D** | ***Niemann-Pick diseases*** |  |  |  |  |
| **63** | *CG3376* | BL 36760 | 29.67 |  |  |
| **64** | *niemann-pick type c-1a* | BL 37504 | 33.66 | BL 38296 | 35.29 |
| **E** | ***Hereditary spastic paraplegias*** |  |  |  |  |
| **65** | *atlastin* | BL 36736 | 30.08 |  |  |
| **66** | *hsp60A* | BL 34729 | 29.15 |  |  |
| **67** | *spichthyin* | BL 44438 | 23.72 | BL 37505 | 27.10 |
| **68** | *spartin* | BL 37499 | 30.22 |  |  |
| **69** | *neuroglian* | BL 37496 | 23.81 | BL 28734 | 23.37 |
| **70** | *M6* | BL 37503 | 33.29 | BL 54032 | 33.87 |
| **71** | *receptor expression enhancing protein A* | BL 37500 | 25.87 |  |  |
| **72** | *seipin* | BL 37501 | 31.44 |  |  |
| **73** | *spastin* | BL 53331 | 26.02 | BL 27570 | 34.81 |
| **74** | *kinesin heavy chain* | BL 25898 | 17.40 | BL 35770,35409 | lethal |
| **75** | *paraplegin* | BL 31100 | 33.78 | BL 31223 | 32.05 |
| **F** | ***Cardiovascular diseases*** |  |  |  |  |
| **76** | *zipper* | BL 36727 | 26.08 | BL 38259,37480 | lethal |
| **77** | *midline* | BL 50681 | 21.66 | BL 38259 | lethal |
| **78** | *tropomyosin 2* | BL 31535 | 18.77 | BL 41695 | 21.29 |
| **79** | *eyes absent* | BL 28733 | 23.82 | BL 67853 | 28.23 |
| **80** | *upheld* | BL 32949 | 20.66 | BL 31541 | lethal |
| **81** | *tropomyosin 1* | BL 56869 | 17.80 | BL 43542 | lethal |
| **82** | *Z band alternatively spliced PDZ-motif* | BL 31561 | 24.52 | BL 58198 | 28.19 |
| **83** | *wings up A* | BL 31893 | 24.82 |  |  |
| **84** | *stress-sensitive B* | BL 31077 | 25.04 | BL 31230 | 21.80 |
| **85** | *dystrophin* | BL 55641 | 21.32 | BL 31553 | 29.85 |
| **86** | *nitric-oxide synthase* | BL 50675 | 23.24 | BL 28792 | 35.55 |
| **87** | *ankyrin* | BL 43965 | 28.92 | BL 31115 | 32.72 |
| **88** | *ankyrin 2* | BL 29438 | 16.25 | BL 33414 | lethal |
| **89** | *angiotensin converting enzyme* | BL 51394 | 29.81 | BL 36749 | lethal |
| **90** | *SNF4/AMP-activated protein kinase gamma* | BL 34726 | 20.53 | BL 26291 | 35.39 |
| **91** | *drosocross 2* | BL 44087 | lethal |  |  |
| **92** | *lamin* | BL 57501 | 25.94 | BL 31605 | 30.42 |
| **93** | *spaghetti-squash activator* | BL 63618 | 26.01 | BL 26735 | 29.35 |
| **94** | *sarcoglycan delta* | BL 55325 | 26.06 | BL 25964 | 25.02 |
| **95** | *drosocross 3* | BL 62456 | 35.98 | BL 62493 | 31.33 |
| **96** | *vinculin* | BL 41959 | 19.08 | BL 25965 | 22.39 |
| **97** | *cheerio* | BL 26307 | 22.22 |  |  |
| **98** | *KCNQ potassium channel* | BL 27252 | 24.16 |  |  |
| **99** | *myosin alkali light chain 1* | BL 27547 | 20.85 |  |  |
| **100** | *sallimus* | BL 31539 | 13.82 | BL 31538 | 20.38 |
| **101** | *CG3803* | BL 31072 | 41.22 |  |  |
| **102** | *Muscle LIM protein at 84B* | BL 31558 | 19.83 |  |  |
| **103** | *myosin light chain 2* | BL 31544 | 21.20 | BL 31543 | 25.30 |
| **104** | *upheld* | BL 32949 | 20.66 | BL 31541 | lethal |
| **105** | *tafazzin* | BL 31694 | 24.01 | BL 31099 | 25.96 |
| **106** | *drosocross1* | BL 31931 | 25.87 |  |  |
| **107** | *hand* | BL 28977 | 24.35 |  |  |
| **108** | *tinman* | BL 28539 | 19.82 | BL 50663 | 18.79 |
| **109** | *actin 57B* | BL 31551 | 26.41 |  |  |
| **110** | *alpha actinin* | BL 34874 | lethal |  |  |
| **111** | *troponin C at 41C* | BL 27053 | 25.71 |  |  |
| **112** | *G protein beta subunit 13F* | BL 35041 | 34.98 | BL 31134 | 36.18 |
| **113** | *lamin C* | BL 31621 | 27.54 |  |  |
| **114** | *G protein beta subunit 13F* | BL 35041 | 34.98 | BL 31134 | 36.18 |
| **115** | *myosin heavy chain* | BL 26299 | lethal | BL 35729 | lethal |
| **116** | *muscle LIM protein at 60A* | BL 29381 | 23.71 |  |  |
| **117** | *pannier* | BL 34659 | 17.99 | BL 33744 | 25.54 |
| **118** | *cac* | BL 27244 | 24.80 |  |  |
| **119** | *actin 87E* | BL 42652 | lethal |  |  |
| **120** | *troponin C at 47D* | BL 26172 | 26.54 | BL 64003 | 33.35 |
| **G** | ***Non-syndromic X-linked intellectual disability*** |  |  |  |  |
| **121** | *GluRIIA* | BL 40907 | 28.94 | BL 27521 | 32.38 |
| **122** | *rab39* | BL 53995 | 31.74 | BL 25953 | 31.64 |
| **123** | *acyl-CoA synthetase long-chain* | BL 27729 | 28.52 | BL 41885,43268 | lethal |
| **124** | *p-21 activated kinase* | BL 62201 | 20.60 | BL 41714 | 28.36 |
| **125** | *BRWD3* | BL 33421 | 11.46 |  |  |
| **H** | ***Alzheimer diseases*** |  |  |  |  |
| **126** | *scully* | BL 41884 | 25.30 | BL 42476 | 26.85 |
| **127** | *tau* | BL 40875 | 16.53 | BL 28891 | 25.52 |
| **128** | *presenilin* | BL 27681 | 24.35 | BL 38374 |  |
| **129** | *nicastrin* | BL 57497 | 20.54 | BL 27498 | 31.38 |
| **130** | *anterior pharynx defective 1* | BL 38249 | 31.23 |  |  |
| **131** | *beta- amyloid protein precursor like* | BL 28043 | 25.00 | BL 39013 | 29.26 |
| **132** | *presenilin enhancer* | BL 52908 | 27.97 | BL 27298 | 33.70 |
| **133** | *par-1* | BL 32410 | 26.35 | BL 35342 | 27.45 |
| **I** | ***Angelman syndrome*** |  |  |  |  |
| **134** | *ubiquitin protein ligase E3A* | BL 31972 | 27.11 | BL 57151 | 35.37 |
| **J** | ***Spinocerebellar ataxia type-2*** |  |  |  |  |
| **135** | *ataxin-2* | BL 67878 | 22.47 | BL 44012 | 29.51 |
| **K** | ***Spinal muscular atrophy*** |  |  |  |  |
| **136** | *survival motor neuron* | BL 67950 | 14.39 | BL 26288 | 21.39 |
| **L** | ***Glycerol kinase deficiency*** |  |  |  |  |
| **137** | *glycerol kinase 2* | BL 57522 | 23.12 | BL 35164 | 33.80 |
| **138** | *glycerol kinase 1* | BL 35188 | 25.41 | BL 51849 | 31.00 |
| **M** | ***Primary microcephaly*** |  |  |  |  |
| **139** | *wd repeat domain 62* | BL 53242 | 33.46 |  |  |
| **140** | *asterless* | BL 38220 | 27.91 | BL 35039 | 29.45 |
| **141** | *microcephalin* | BL 38244 | 33.32 |  |  |
| **142** | *abnormal spindle* | BL 28741 | 25.84 | BL 35224 | 37.30 |
| **143** | *centrosomin* | BL 31402 | 24.11 | BL 35761 | 38.84 |
| **144** | *spindle assembly abnormal 4* | BL 35049 | 36.16 |  |  |
| **N** | ***Peroxisome biogenesis disorders-Zellweger spectrum*** |  |  |  |  |
| **145** | *peroxin 3* | BL 50694 | lethal |  |  |
| **146** | *alkylglycerone phosphate synthase* | BL 34350 | 32.18 |  |  |
| **147** | *peroxin 13* | BL 50697 | 25.61 |  |  |
| **148** | *peroxin 12* | BL 53308 | 26.53 |  |  |
| **149** | *peroxin 16* | BL 57495 | 29.06 |  |  |
| **150** | *peroxin 11* | BL 42883 | 30.72 |  |  |
| **151** | *peroxin 19* | BL 50702 | 28.53 |  |  |
| **152** | *peroxin 5* | BL 42854 | 20.49 | BL 58064 | 36.77 |
| **153** | *CG2316* | BL 41984 | 24.86 |  |  |
| **154** | *peroxin 1* | BL 28979 | 23.49 | BL 51497 | 29.20 |
| **O** | ***P-type ATPase disorders*** |  |  |  |  |
| **155** | *secretory pathway calcium ATPase* | BL 44040 | 35.60 | BL 28352 | 33.09 |
| **156** | *sarco/endoplasmic reticulum Ca(2+)-ATPase* | BL 25928 | 17.06 | BL 44581 | 30.33 |
| **157** | *Na(+)/K(+)-exchanging ATPase* | BL 32913 | 35.90 | BL 28073 | lethal |
| **P** | ***Cornelia de Lange syndrome*** |  |  |  |  |
| **158** | *verthandi* | BL 36786 | 21.21 |  |  |
| **159** | *structural maintenance of chromosomes 1* | BL 34351 | 31.81 |  |  |
| **160** | *structural maintenance of chromosomes 3* | BL 33431 | 21.19 | BL 50899 | 30.94 |
| **161** | *nipped-B* | BL 32406 | 18.98 | BL 28961 | 30.75 |
| **Q** | ***Neuronal ceroid lipofuscinosis*** |  |  |  |  |
| **162** | *palmitosyl-protein thioesterase 1* | BL 62291 | 37.65 | BL 25952 | 30.75 |
| **163** | *cln3* | BL 35734 | 30.87 |  |  |
| **164** | *cathepsin D* | BL 53882 | 35.41 | BL 55178 | 29.88 |
| **R** | ***Werner syndrome*** |  |  |  |  |
| **165** | *wrn exonuclease* | BL 38297 | 32.48 |  |  |
| **S** | ***Hypoparathyroidism-retardation-dysmorphism syndrome*** |  |  |  |  |
| **166** | *tubulin-specific chaperone E* | BL 44569 | 21.80 |  |  |
| **T** | ***Amylotrophic lateral sclerosis*** |  |  |  |  |
| **167** | *TAR DNA-binding protein-43 homolog* | BL 39014 | 20.01 | BL 29517 | 28.07 |
| **168** | *5-hydroxytryptamine receptor 1B* | BL 25833 | 26.88 |  |  |
| **169** | *FIG4 phosphoinositide 5-phosphatase* | BL 58063 | 26.03 | BL 38291 | 27.45 |
| **170** | *vamp- associated protein 33kDa* | BL 27312 | 28.69 |  |  |
| **171** | *sod1* | BL 29389 | 21.33 | BL 34616 | 25.76 |
| **172** | *cabeza* | BL 34839 | 25.38 | BL 32990 | 26.93 |
| **U** | ***Huntington diseases*** |  |  |  |  |
| **173** | *huntingtin* | BL 31264 | 30.66 | BL 44550 | 30.45 |
| **174** | *huntingtin-interacting protein 14* | BL 35012 | 19.15 | BL 31591 | 24.90 |
| **V** | ***Neurofibromatosis type 1*** |  |  |  |  |
| **175** | *neurofibromin 1* | BL 31466 | 26.86 | BL 25845, 53322 | lethal |
| **W** | ***Fragile X-syndrome*** |  |  |  |  |
| **176** | *fmr1* | BL 34944 | 24.94 | BL 27484 | 29.69 |
| **X** | ***Coffin-Lowry syndrome*** |  |  |  |  |
| **177** | *ribosomal protein S6 kinase II* | BL 27731 | 32.62 | BL 56031 | 29.43 |
| **Y** | ***X-linked infantile spinal muscular atrophy*** |  |  |  |  |
| **178** | *ubiquitin activating enzyme 1* | BL 36307 | 12.77 | BL 25957 | 25.94 |
| **Z** | ***Peroxisome disorders ( non-Zellweger spectrum)*** |  |  |  |  |
| **179** | *acyl-coA oxidase 1* | BL 52882 | 28.42 |  |  |
| **AB** | ***ER-stress response factors*** |  |  |  |  |
| **180** | *atf6* | BL 26211 | 39.49 |  |  |
| **181** | *Inositol-requiring enzyme-1* | BL 35253 |  |  |  |
| **182** | *pancreatic elF-2 alpha kinase* | BL 42499 |  |  |  |
| **183** | *X-box binding protein-1* | BL 25990 |  |  |  |
| **184** | *chop24* | BL 64563 |  |  |  |
| **185** | *CG11857* | BL 57435 | 25.09 |  |  |
| **AC** | ***PolyQ disorders*** |  |  |  |  |
| **186** | *UAS-HTT128Q.FL* | BL 33808 | 38.25 |  |  |
| **187** | *UAS-CAG.20Q* | BL 30549 | 31.42 |  |  |
| **188** | *UAS-Q108* | BL 68393 | 35.23 |  |  |
| **189** | *UAS-41Q.HA* | BL 30540 | 34.73 |  |  |
| **190** | *GluRIII RNAi* (Control) |  | 35.22 |  |  |

**Table S1:** Table showing the source of *Drosophila* stocks, gene name and mean EPSP amplitudes of 300 RNAi lines homologs with human neurological and muscle-related disorder links. BL stands for Bloomington stock number and Sr. No. denotes serial number, to count table rows.
