## Supplementary material for "Roles for Mitochondrial Complex I subunits in regulating synaptic transmission and growth": Table S2

| **Sr. No.** | **Gene name** | **pre+post-Gal4 or (C15) (mEPSP,EPSP in mV,** **Freq. in Hz)** | ***p*-value**  **(EPSP)**  **Control vs. C15>RNAi** | **Pre+post-Gal4+ *GluRIII* RNAi or (T15)**  **(mEPSP,EPSP in mV, Freq. in Hz)** | ***p*-value**  **(EPSP)**  **C15>RNAi vs. T15>RNAi** |
| --- | --- | --- | --- | --- | --- |
| **1** | Control | mEPSP (0.75 ± 0.03), EPSP (37.67 ± 2.19), QC (49.70 ± 1.71), Freq. (3.01 ± 0.30), n=8 |  | mEPSP (0.53 ± 0.03), EPSP (38.41 ± 2.20), QC (77.30 ± 2.74), Freq. (1.96 ± 0.30), n=6 | p=0.8208 |
| **2** | *NDUFS1 (CG2286/*  *ND-75)* | mEPSP (0.54 ± 0.02), EPSP (19.30 ± 1.50), QC (32.54 ± 0.31), Freq. (4.17 ± 1.26), n=9 | p< 0.0001 | mEPSP (0.36 ± 0.02), EPSP (16.53 ± 0.88), QC (47.17 ± 3.58), Freq. (2.73 ± 0.71), n=13 | p=0.1058 |
| **3** | *NDUFS2 (CG1970/*  *ND-49)* | mEPSP (0.59 ± 0.04), EPSP (24.66 ± 1.50), QC(42.57 ± 2.47), Freq. (3.86 ± 0.29), n=8 | p=0.0002 | mEPSP (0.40 ± 0.02), EPSP (23.71 ± 0.65), QC (60.18 ± 3.87), Freq. (1.46 ± 0.18), n=8 | p=0.5712 |
| **4** | *NDUFS3 (CG12079/*  *ND-30)* | mEPSP (0.53 ± 0.03), EPSP (26.96 ± 1.22), QC (50.76 ± 2.21, Freq.(3.76 ± 0.58), ns, n=6 | p=0.0022 | mEPSP (0.43 ± 0.02), EPSP (31.17 ± 0.93), QC (72.85 ± 4.06), Freq. ( 1.247 ± 0.13), n=6 | p=0.0210 |
| **5** | *NDUFS7 (CG9172/ND-20)* | mEPSP (0.72 ± 0.07), EPSP (26.29 ± 1.41), QC (38.52 ± 4.54), Freq. (3.42 ± 0.23), n=7 | p= 0.0010 | mEPSP (0.57 ± 0.02), EPSP (27.71 ± 1.27), QC (48.97 ± 3.44), Freq. ( 0.95± 0.10), n=7 | p=0.4708 |
| **6** | *NDUFS7 (CG2014/ND-20L)* | mEPSP (0.80 ± 0.03), EPSP (27.44 ± 0.70), QC (34.44 ± 1.34), Freq. (6.46 ± 0.41), n=8 | p=0.0006 | mEPSP (0.44 ± 0.05), EPSP (19.55 ± 0.68), QC (48.18 ± 6.95), Freq. (3.86 ± 1.17), n=6 | p< 0.0001 |
| **7** | *NDUFS8 (CG3944/*  *ND-23)* | mEPSP (0.67 ± 0.03), EPSP (29.50 ± 1.54), QC (44.32 ± 3.45), Freq. (3.38 ± 0.29), n=6 | p=0.0149 | mEPSP (0.44 ± 0.02), EPSP (26.96 ± 1.04), QC (60.81 ± 3.00), Freq.(2.95 ± 1.68), n=6 | p=0.2021 |
| **8** | *NDUFV1 (CG9140/ND-51)* | mEPSP (0.39 ± 0.05), EPSP (14.03 ± 1.21), QC ( 65.76 ± 3.07), Freq . (6.32 ± 2.29), n=6 | p< 0.0001 | mEPSP (0.27 ± 0.01), EPSP (15.83 ± 1.13), QC ( 88.23 ± 11.43), Freq. (2.40 ± 1.48), n=6 | p=0.3034 |
| **9** | *NDUFV1 (CG11423/ND51L1)* | mEPSP (0.55 ± 0.03), EPSP (32.07 ± 1.37), QC ( 59.75 ± 6.16), Freq. (2.91 ± 0.47), n=6 | p=0.0699 | mEPSP (0.40 ± 0.02), EPSP (30.89 ± 2.19), QC ( 78.93 ± 6.80), Freq. ( 1.24± 0.12), n=8 | p=0.6823 |
| **10** | *NDUFV1 (CG8102/ND-51L2)* | mEPSP (0.65 ± 0.03), EPSP (24.01 ± 1.14), QC (37.77 ± 2.33), Freq. (3.37 ± 0.15), n=7 | p=0.0001 | mEPSP (0.40 ± 0.02), EPSP (22.08 ± 1.77), QC (55.08 ± 4.00), Freq. (1.484 ± 0.11), n=11 | p=0.4374 |
| **11** | *NDUFV2 (CG5703/ND-24)* | mEPSP (0.78 ± 0.04), EPSP (27.86 ± 3.28), QC ( 36.33 ± 4.46), Freq. (3.78 ± 0.49), n=8 | p=0.0262 | mEPSP (0.45 ± 0.02),EPSP (26.38 ± 3.26), QC ( 57.96 ± 6.11), Freq. (0.36 ± 0.08), n=6 | p=0.7602 |
| **12** | *NDUFV2 (CG6485/ND-24L*) | mEPSP (0.61 ± 0.06), EPSP (32.95 ± 1.96), QC ( 56.25 ± 6.42), Freq. (2.83 ± 0.35), n=6 | p= 0.1489 | mEPSP (0.36 ± 0.01), EPSP (29.42 ± 2.62), QC ( 80.43 ± 6.78), Freq. (1.53 ± 0.78), n=7 | p=0.3173 |
| **13** | *ND-1 (CG34092/mt:ND-1)* | mEPSP (0.66 ± 0.05), EPSP (32.98 ± 1.62), QC ( 50.57 ± 3.33). Freq. (4.35 ± 0.47), n=7 | p=0.1174 | mEPSP (0.46 ± 0.01), EPSP (30.38 ± 2.99), QC ( 66.44 ± 7.68), Freq. (2.060 ± 0.19), n=7 | p=0.4592 |
| **14** | *ND-2 (CG34063/mt:ND-2)* | mEPSP (0.69 ± 0.02), EPSP (32.15 ± 1.45), QC ( 51.33 ± 3.34), Freq. (1.74 ± 0.14), n=7 | p=0.0630 | mEPSP (0.35 ± 0.02), EPSP (23.12 ± 2.40), QC ( 84.70 ± 4.76), Feeq. (1.53 ± 1.07), n=9 | p=0.0099 |
| **15** | *ND-3 (CG34076/mt:ND-3)* | mEPSP (0.69 ± 0.04),EPSP (31.88 ± 2.35),  QC ( 46.57 ± 3.93), Freq. (5.85 ± 1.38), n=8 | p=0.0934 | mEPSP (0.45 ± 0.02), EPSP (24.34 ± 2.11), QC ( 53.88 ± 3.93), Freq. (1.13 ± 0.11), n=9 | p=0.0305 |
| **16** | *ND-4 (CG34058/mt:ND-4)* | mEPSP (0.79 ± 0.03), EPSP (32.42 ± 3.03), QC ( 41.80 ± 4.97), Freq. (2.19 ± 0.41), n=6 | p=0.1751 | mEPSP (0.34 ± 0.02), EPSP (29.91 ± 3.0), QC ( 77.37 ± 9.54), Freq. ( 0.75 ± 0.16), n=9 | p=0.5737 |
| **17** | *ND-4L (CG34086/*  *mt:ND-4L)* | mEPSP (0.52 ± 0.02), EPSP (25.33 ± 1.74), QC (48.99 ± 3.54), Freq. (3.32 ± 0.35), n=7 | p= 0.0008 | mEPSP (0.37 ± 0.01), EPSP (20.10 ± 1.23), QC (53.60 ± 4.23), Freq. (1.23 ± 0.12), n=9 | p=0.0242 |
| **18** | *ND-5 (CG34083/*  *mt:ND-5)* | mEPSP (0.67 ± 0.05), EPSP (22.54 ± 2.47), QC (34.57 ± 4.27), Freq. (7.10 ± 0.48), n=7 | p=0.0004 | mEPSP (0.43 ± 0.02), EPSP (25.49 ± 1.94), QC (55.91 ± 7.24), Freq. (1.55 ± 0.26), n=6 | p=0.3633 |
| **19** | *ND-6 (CG34089/*  *mt:ND-6)* | mEPSP (0.72 ± 0.03), EPSP (23.63 ± 1.57), QC (32.76 ± 2.42), Freq. (3.40 ± 0.35), n=7 | p=0.0002 | mEPSP (0.49 ± 0.03), EPSP (25.28 ± 1.02), QC (52.57 ± 3.67), Freq. (1.47 ± 0.21), n=11 | p=0.3703 |

**Table S2:** This table shows mEPSP, EPSP amplitudes, Frequency and calculated QC of 11 nuclear- and 7 mitochondrial-encoded complex I subunits in pre- and post-Gal4 driven CI RNAi and CI RNAi along with *GluRIII* RNAi. *ND-1*, *ND-2*, *ND-3*, *ND-4*, *ND-4L*, *ND-5* and *ND-6* represent mitochondrial encoded complex I subunits. *p-*values are described in the table for each analysis. Error represents mean±s.e.m. Statistical analysis based on one-way ANOVA with Tukey’s post-hoc vs. the control data set at the top of the column. Student’s t-test, comparing evoked amplitudes (EPSP) in column 3 vs. column 5. Sr. No. stands for serial number. “C15” and “T15” are not full genotypes. They are shorthand for screening lines containing pre- and postsynaptic Gal4 drivers (C15 and T15) and *GluRIII[RNAi]* (T15) (Brusich et al., 2015).
