## Supplementary material for "Roles for Mitochondrial Complex I subunits in regulating synaptic transmission and growth": Table S3

| **Sr. No.** | **Genotypes** | **mEPSP amplitude (mV)** | **EPSP amplitude (mV)** | **mEPSP Freq. (Hz)** | **Quantal content (QC)** | **Distance**  **Crawled**  **(in cm)** |
| --- | --- | --- | --- | --- | --- | --- |
| **1** | *elaV-(C155)-Gal4/+*, | 0.70 ± 0.03, n*=*6 | 38.99 ± 2.35, n=6 | 2.55 ± 0.46, n=6 | 55.22 ± 2.76, n=6 | 2.07 ± 0.17, n=14 |
| **2** | *elaV-(C155)/+; ND-20L*  *RNAi/+* | 0.75 ± 0.04, n=7 | 37.99 ± 1.44, n=7 | 5.41 ± 0.55, n=7 | 50.51 ± 1.94, n=7 | 2.51 ± 0.15, n=14 |
| **3** | *BG-57-Gal4/+* | 0.89 ± 0.03, n=7 | 42.62 ± 1.18, n=7 | 3.47 ± 0.50, n=7 | 48.13 ± 2.36, n=7 | 2.72 ± 0.18, n=15 |
| **4** | *BG-57-Gal4/+; ND-20L RNAi/+* | 0.65 ± 0.03, n=10 | 27.49 ± 1.20, n=10 | 5.39 ± 0.60, n=10 | 43.07 ± 2.55, n=10 | 0.64 ± 0.08, n=11 |

**Table S3:** This table shows mEPSP, EPSP amplitudes in (mV), Frequency, calculated QC and crawling behavior of the genotypes as indicated in the table. Muscle depleted *ND-20L* RNAi showed reduced EPSP amplitudes and crawling ability, while pan-neuronal *ND-20L* larvae did not show any remarkable phenotypes. Error represents mean±s.e.m. Statistical analysis based on Student’s t-test for pairwise comparison. Sr. No. denotes serial number.
