## Supplementary material for "Roles for Mitochondrial Complex I subunits in regulating synaptic transmission and growth": Table S4

| **Sr.**  **No.** | **Condition** | **mEPSP, EPSP in mV and Freq. in Hz (Respective Control)** | **Condition** | **mEPSP, EPSP in mV and Freq. in Hz (Rotenone)** | **p-value**  **(EPSP)**  **Control vs. Drug** |
| --- | --- | --- | --- | --- | --- |
| **1** | ***w^1118^*^,^ 5% DMSO, 48 hrs** | mEPSP (0.86 ± 0.02), EPSP (40.37 ± 1.78), QC (48.04 ± 3.10), Freq. (5.58 ± 0.53), n=7 | ***w^1118^*, 2µM rotenone, 48 hrs** | mEPSP (1.11 ± 0.03), EPSP (40.38 ± 0.84), QC (36.40 ± 1.36), Freq. (4.85 ± 0.64), n=6 | p=0.9978 |
| **2** | ***w^1118^*, 5% DMSO, 30 minutes open prep** | mEPSP (0.67 ± 0.08), EPSP (35.76 ± 1.35), QC (57.33 ± 6.42), Freq. (4.79 ± 0.62), n=7 | ***w^1118^*, 500µM rotenone, 30 minutes open prep** | mEPSP (0.68 ± 0.03), EPSP (31.91 ± 0.72), QC (47.01 ± 2.85), Freq. (5.15 ± 0.44), n=6 | p=0.0275 |
| **3** | ***w^1118^*, 5% DMSO, 72 hrs** | mEPSP (0.94 ± 0.03), EPSP (43.25 ± 1.33), QC (46.21 ± 2.49), Freq. (4.53 ± 0.47), n=6 | ***w^1118^*, 500µM rotenone, 72 hrs** | mEPSP (1.24 ± 0.05), EPSP (34.70 ± 2.10), QC (28.45 ± 2.74), Freq. (7.52 ± 0.42), n=6 | p=0.0064 |
| **4** | ***w^1118^*, 5% DMSO, 7 hours liquid food** | mEPSP (0.91 ± 0.08), EPSP (40.87 ± 0.86), QC (46.86 ± 4.22), Freq. (6.17 ± 0.99), n=7 | ***w^1118^*, 500µM rotenone, 7 hrs liquid food** | mEPSP (0.73 ± 0.06), EPSP (33.17 ± 0.75), QC (48.48 ± 3.92), Freq. (6.39 ± 1.30), n=7 | p< 0.0001 |
| **5** | ***w^1118^*, 0.5% DMSO, embryo to third instar** | mEPSP (0.92 ± 0.06), EPSP (38.09 ± 1.25), QC (42.46 ± 2.84), Feeq. (7.01 ± 1.04), n=9 | ***w^1118^*, 25µM rotenone, embryo to third instar** | mEPSP (0.82 ± 0.05), EPSP (30.52 ± 1.62), QC (38.86 ± 3.66), Freq. (4.40 ± 0.33), n=9 | p=0.0020 |
| **6** | ***w^1118^*, 1% DMSO, embryo to third instar** | mEPSP (0.91 ± 0.03), EPSP (34.78 ± 1.11), QC (38.10 ± 1.31), Freq. (4.84 ± 0.56), n=8 | ***w^1118^*, 50µM rotenone, embryo to third instar** | mEPSP (0.61 ± 0.02), EPSP (25.30 ± 1.35), QC (41.74 ± 2.70), Freq. (4.81 ± 0.37), n=9 | p< 0.0001 |

**Table S4**: This table shows mEPSP and EPSP amplitudes (mV), mEPSP frequency (Hz) and QC analysis of third instar wild type-larvae treated or fed either with DMSO (carrier, column 2) or rotenone + DMSO at various concentrations (column 4). Rotenone-treated larvae affect evoked vesicle release of neurotransmitters compared to the control animals. Error represents mean±s.e.m. p-values are represented in column 5 for each analysis. Statistical analysis based on Student’s t-test, comparing experimental condition (rotenone) vs. control condition (DMSO carrier). Sr. No. denotes serial number.
