## Supplementary material for "Roles for Mitochondrial Complex I subunits in regulating synaptic transmission and growth": Table S5

| **Sr. No.** | ***Drosophila* gene name , Bloomington stocks, Antibodies, dilution and Fluorophores** |
| --- | --- |
| **1** | *ND-75* RNAi*/CG2286* (BL 33911, BL-27739), *ND-49* RNAi*/CG1970* (BL 28573, BL 57499), *ND-30* RNAi*/CG12079* (BL 44535, BL-51425) |
| **2** | *ND-20* RNAi*/CG9172* (BL 64995*), ND-20L* RNAi*/CG2014* (BL 62381), *ND-23* RNAi*/CG3944* (BL 51797, BL-30487) |
| **3** | *ND-51*RNAi */CG9140* (BL 52939, BL-36701, BL-29534), *ND-51L1* RNAi*/CG11423* (BL 42591), *ND-51L2* RNAi*/CG8102* (BL 64536) |
| **4** | *ND-24* RNAi*/CG5703* (BL 51855), *ND-24L* RNAi*/CG6485* (BL 60873) |
| **5** | *ND-1 RNAi/CG34092* (BL 64852), *ND-2* RNA*i/CG34063* (BL 77154), *ND-3* RNAi*/CG34076* (BL 77153), *ND-4* RNAi*/CG34085* (BL 65216), *ND-4L* RNAi*/CG34086* (BL 65158), *ND-5* RNAi/*CG34083* (BL 77336), *ND-6* RNAi/*CG34089* (BL 64670) |
| **6** | Mouse α-DLG (DSHB-1:50), Mouse α-Synapsin (1:30), Alexa α-HRP 488 (1:800), Mouse Alexa Fluor 568 (1:400), Mouse Alexa Fluor 488 (1:400), Polyclonal rabbit α-DLG (1:700) |

**Table S5:** Table showing source of *Drosophila* stocks, primary antibodies, secondary antibodies and fluorophores. Sr. No. denotes serial number.
